# Phosphosite Scanning reveals a complex phosphorylation code underlying CDK-dependent activation of Hcm1

**DOI:** 10.1101/2022.08.04.502803

**Authors:** Michelle M. Conti, Rui Li, Michelle A. Narváez Ramos, Lihua Julie Zhu, Thomas G. Fazzio, Jennifer A. Benanti

**Affiliations:** Department of Molecular, Cell and Cancer Biology, University of Massachusetts Chan Medical School, Worcester, MA 01605, USA; Program in Bioinformatics and Integrative Biology, University of Massachusetts Chan Medical School, Worcester, MA 01605, USA

## Abstract

Ordered cell cycle progression is coordinated by cyclin dependent kinases (CDKs). CDKs often phosphorylate substrates at multiple sites clustered within disordered regions. However, for most substrates, it is not known which phosphosites are functionally important. We developed a high-throughput approach, Phosphosite Scanning, that tests the importance of each phosphosite within a multisite phosphorylated domain. We show that Phosphosite Scanning identifies multiple combinations of phosphosites that can regulate protein function and reveals specific phosphorylations that are required for phosphorylation at additional sites within a domain. We applied this approach to the yeast transcription factor Hcm1, a conserved regulator of mitotic genes that is critical for accurate chromosome segregation. Phosphosite Scanning revealed a complex CDK-regulatory circuit that mediates processive phosphorylation of key activating sites *in vivo*. These results illuminate the mechanism of Hcm1 activation by CDK and establish Phosphosite Scanning as a powerful tool for decoding multisite phosphorylated domains.

## Introduction

Progression through the cell division cycle requires the coordinated function of hundreds of proteins. This coordination is achieved in part through phosphorylation of numerous substrates by cyclin-dependent kinase (CDK) complexes^1–5^. This critical mechanism of cell cycle control is conserved from yeast to humans^6, 7^. Phosphorylation of CDK substrates can trigger a variety of actions, including substrate degradation or activation, interactions of substrates with other proteins, and re-localization of substrates to different sub-cellular compartments. These changes have widespread implications in cell cycle control, including the regulation of key phase transitions and the expression of cyclically transcribed genes.

A common feature among CDK substrates is that they are often phosphorylated on multiple sites. CDKs are serine and threonine directed kinases that efficiently phosphorylate an optimal consensus motif (S/T-P-x-K/R) *in vitro*^8–10^. However, a minimal consensus motif (S/T-P) is frequently phosphorylated *in vivo*. These consensus motifs often occur in clusters and are enriched in intrinsically disordered regions (IDRs)^3, 11, 12^. Despite these common features, the numbers and/or positions of phosphates required to elicit functional changes vary considerably among CDK substrates. Some substrates contain many CDK phosphorylation sites, but functional changes depend on phosphorylation of only one or a few specific sites.

Phosphorylation at a specific site can be required to trigger structural changes that impact protein function^13–15^, or to generate a linear binding epitope^16, 17^. Conversely, in a different group of substrates, phosphorylation at many sites is required to change the overall charge of a domain and elicit a functional change^18^. For most CDK substrates, the CDK regulatory circuitry remains poorly understood, including (1) the fraction of phosphosites required to regulate protein function and (2) which particular sites, among many that are phosphorylated by CDKs, have regulatory function.

Determining how individual phosphosites contribute to protein regulation has remained a significant challenge because it requires generating large numbers of mutant proteins for any given substrate, and evaluating the effects of each mutant one by one. Although it may be feasible to generate a series of mutants in which each phosphosite is mutated individually, this strategy only examines the importance of each phosphosite in an otherwise wild-type context, potentially missing effects observed when combinations of phosphosites are mutated. One streamlined approach that has been developed to systematically examine the effects of mutations on protein function in a pooled format is deep mutational scanning^19–22^. Most deep mutational scanning approaches analyze all possible amino acid changes at each residue within a domain, which provides high resolution information about protein structure. Here, we modified deep mutational scanning to focus specifically on individual phosphosites within a multiphosphorylated domain, and to determine how phosphorylation at each site impacts protein function *in vivo*. Our modified approach, called Phosphosite Scanning, uses bulk competition assays to measure the cellular fitness outcomes caused by mutation of phosphorylation sites to unphosphorylatable or phosphomimetic residues, in all possible combinations. By comparing the fitness effects of proteins with different combinations of wild type, unphosphorylatable and phosphomimetic residues, it is possible to determine which phosphosites contribute to protein regulation, the combinations of phosphosites that are necessary and sufficient for normal activity, and how phosphorylation at a particular site impacts phosphorylation at additional sites within a domain.

To examine how CDK can regulate protein function through multisite phosphorylation, we applied Phosphosite Scanning to the yeast forkhead family transcription factor Hcm1. Like its human homolog, FoxM1, Hcm1 activates expression of genes that are required for mitotic spindle function and accurate chromosome segregation^23, 24^. Notably, mutations in Hcm1 impact cellular fitness. Mutation of all CDK phosphoacceptor sites in Hcm1 to alanine inactivates the protein and reduces cellular fitness^25^. Conversely, mutation of all eight phosphoacceptor sites within the transactivation domain (TAD) to two glutamic acid residues—mimicking the charge of a phosphate—results in elevated activity and increased fitness compared to wild type^26^. These phosphorylation dependent fitness effects make the Hcm1 TAD an ideal model to investigate the mechanism of phosphoregulation by CDK.

Phosphosite Scanning of Hcm1 revealed several features of activation that highlight the complex regulation of multisite phosphorylated domains. Remarkably, we found that non-overlapping combinations of phosphomimetic mutations were able to activate Hcm1. Phosphomimetic mutations at two central phosphosites, T460 and S471, had the greatest impact on Hcm1 activity and, on their own, could restore wild type activity when all other phosphosites were changed to alanine. However, phosphomimetic mutations at all sites except T460 and S471 led to similar activation, indicating a mechanism in which both the position and number of CDK sites are important for activation. We also found that phosphorylation of three N-terminal threonine residues within the TAD (T428, T440, and T447) was necessary for phosphorylation of T460 and S471. Our results suggest that CDK sites within the TAD are phosphorylated processively via the CDK phosphoadaptor subunit Cks1, which interacts specifically with phosphothreonines to promote increased phosphorylation. Notably, these insights were only made possible by the ability of Phosphosite Scanning to interrogate all possible combinations of phosphodead and phosphomimetic mutations, as well as the ability to quantify the effects of mutations in multiple contexts (i.e., each combination of phosphomutants is assayed in the presence of wild type or mutant configurations of all other phosphosites within the domain). In sum, our results reveal the mechanism of Hcm1 activation by CDK and establish Phosphosite Scanning as a powerful approach that can be used to interrogate multisite phosphorylated domains.

## Results

### Phosphorylation of the Hcm1 TAD regulates fitness

Hcm1 regulation by CDK is complex: it contains 11 minimal CDK consensus sites that regulate both its degradation and transcriptional activity^25^. Like other CDK substrates, most of these sites fall within IDRs (Fig. 1a). A cluster of three sites in the Hcm1 N-terminus constitutes a phosphodegron that, when phosphorylated, signals for its degradation by an SCF-family ubiquitin ligase^25^. A second cluster of eight sites is located within the TAD and is required for Hcm1 to activate transcription of target genes. However, the specific requirements for phosphoregulation of the TAD are not understood.

**Fig. 1.**
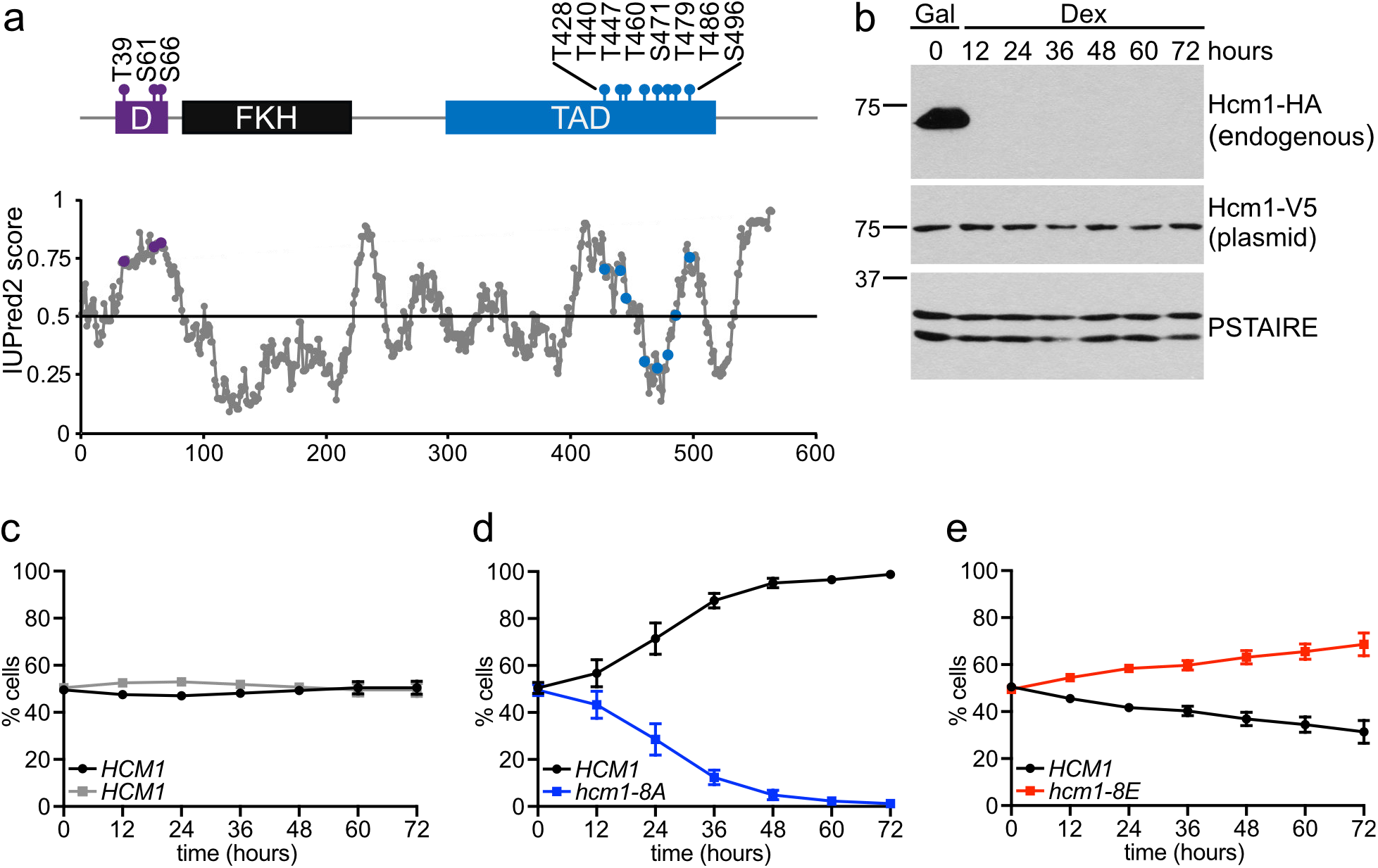
Phosphorylation of the Hcm1 TAD regulates fitness. **(a)** Schematic showing previously characterized CDK consensus sites (S/T-P), characterized domains and predicted disordered regions. D indicates the phosphodegron, FKH indicates the forkhead DNA binding domain, TAD indicates the transcriptional activation domain. IUPred2 disorder prediction shown with phosphorylation sites in purple (D) and blue (TAD). **(b)** Western blot showing the expression of the indicated proteins at the timepoints shown. Strains were initially grown in galactose (Gal) to allow expression of the 3HA-tagged wild type protein from the genome and were shifted to dextrose (Dex) to repress wild type expression at the start of the experiment. Plasmid expressed Hcm1 proteins were detected with an antibody that recognizes a C-terminal 3V5 tag, PSTAIRE is shown as a loading control. **(c-e)** Strains expressing the indicated *HCM1* alleles from plasmids were co-cultured and the percentage of cells expressing each allele was determined at the time points indicated. Shown is an average of n = 3 experiments, error bars represent standard deviations.

Previously, mutation of all 15 CDK phosphoacceptor sites in Hcm1 to alanine was shown to reduce cellular fitness in a pairwise, co-culture assay^25^. To investigate the effects of blocking phosphorylation exclusively within the TAD, we mutated the eight sites in this region to alanine (*hcm1-8A*) and assayed the fitness of this strain compared to wild type in a co-culture assay. Low copy centromeric plasmids expressing wild type *HCM1* or *hcm1-8A* from its endogenous promoter were transformed into strains expressing either mutant (non-fluorescent) or wild type GFP, respectively, and in which the galactose-inducible *GAL1* promoter was integrated upstream of the genomic copy of *HCM1*. At the start of the assay, equivalent amounts of each strain were diluted into dextrose-containing medium to shut off expression of endogenous *HCM1* (Fig. 1b). Co-cultures were then sampled and diluted every 12 hours for a total of 72 hours to prevent cells from reaching saturation and exiting the cell cycle. At the end of the experiment, the percentage of each strain in the population at each time point was quantified by flow cytometry. As a control, co-culture assays with wild type strains confirmed equal fitness of strains expressing functional or mutant GFP markers (Fig. 1c). Consistent with a role for TAD phosphorylation in Hcm1 function, *hcm1-8A* cells were dramatically less fit than wild type cells (Fig. 1d). Conversely, a gain of function allele that mimics full phosphorylation of the TAD through mutation of all S/T-P sites to two glutamic acid residues (*hcm1-8E*) conferred a fitness benefit when compared to wild type, as previously reported (previously referred to as *hcm1-16E*^26^) (Figure 1E). We sought to leverage these opposing effects on fitness to investigate the contribution of each CDK phosphorylation site within the TAD to Hcm1 function.

### Phosphosite Scanning: decoding Hcm1 phosphoregulation through pooled fitness assays

We developed an approach, called Phosphosite Scanning, in which bulk fitness screens are used to interrogate all possible combinations of phosphorylated sites within a multiphosphorylated domain. In contrast to saturating mutagenesis approaches^19^, this method focuses on amino acid substitutions that mimic or prevent phosphorylation. The fitness of strains expressing phosphomutant proteins can be measured by monitoring the frequency of all mutants in the population over time using deep sequencing. By using fitness as a proxy for protein activity, the impact of phosphorylation at each site on Hcm1 activity can be quantified. Screens utilizing only unphosphorylatable and phosphomimetic mutations quantify the activity of proteins with a fixed phosphorylation state. Additionally, the inclusion of wild type sites along with phosphosite mutations allows us to infer which sites are phosphorylated *in vivo* (described below).

We first designed a set of phosphomutant alleles such that each S/T-P site was mutated to either unphosphorylatable A-P or phosphomimetic E-E, in all possible combinations, to generate a pooled library of 256 distinct phosphomutants (referred to as the A/E library, Fig. 2a). Gene fragments containing phosphosite mutations were assembled by annealing and extending a mixture of overlapping mutagenic oligonucleotides that cover all eight CDK sites within the TAD (Fig. S1a). The fragments were then ligated and cloned into a low copy expression vector containing the *HCM1* promoter. Despite variation in codon usage, phosphosite mutations did not affect Hcm1 protein levels (Extended Data Fig. 1b). The A/E library, which included a construct expressing the wild type gene, was then transformed into a *GAL1p-HCM1* strain in galactose containing medium, similar to the pairwise co-culture assay described above (Fig. 2b). This ensured that wild type Hcm1 was present in cells prior to the start of the screen to prevent selection against strains expressing non-functional or lowly functional *HCM1* mutants. The population was sampled at the start of the screen and cells were then shifted to dextrose containing medium to shut off endogenous *HCM1* expression. The culture was sampled and diluted every 12 hours for a total of 72 hours to prevent the cells from reaching saturation and exiting the cell cycle. The abundance of each Hcm1 mutant in the population at each timepoint was then determined by Illumina sequencing. The screen was performed in triplicate, and replicates were highly correlated (Extended Data Fig. 2a).

**Fig. 2.**
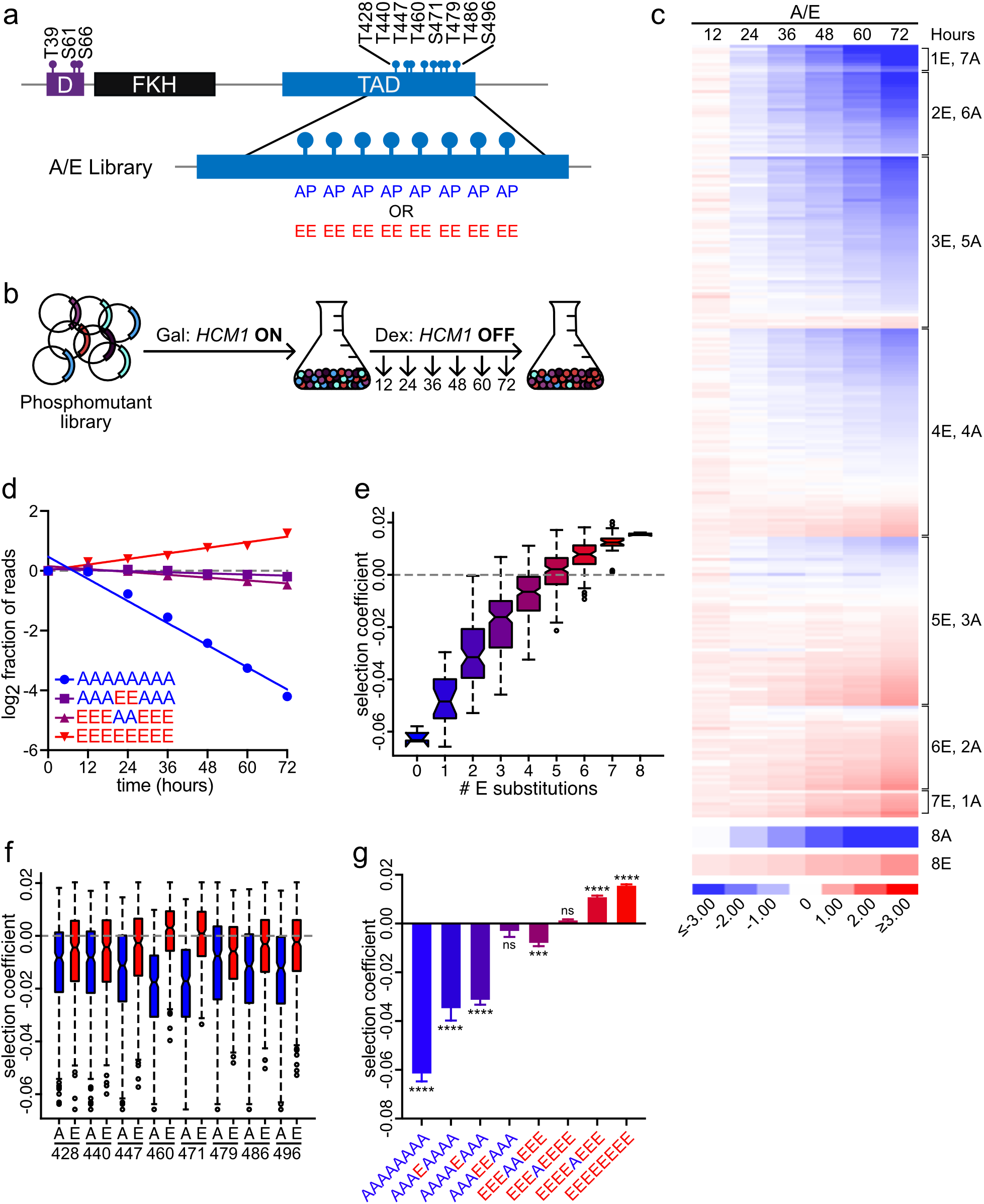
Decoding Hcm1 phosphoregulation with Phosphosite Scanning. **(a)** Schematic of phosphomutant library design. Each S/T-P site within the TAD was mutated to either unphosphorylatable A-P or phosphomimetic E-E in all 256 possible combinations. **(b)** Schematic of the Phosphosite Scanning approach. The plasmid library is transformed into a *GAL1p-HCM1* strain. Cells are shifted from galactose (Gal) to dextrose (Dex) at the start of the screen to repress expression of the wild type protein. The population is sampled and diluted every 12 hours for a total of 72 hours. Mutant abundance at each time point is determined by Illumina sequencing. **(c)** Heat map representation the A/E Phosphosite Scanning screen. Each row represents a mutant, shown is the log2 fold change in normalized read counts with respect to time zero for each mutant, all mutants have been normalized to wild type. Blue indicates depletion, red indicates enrichment. Shown is an average of n = 3 biological replicates. See Extended Data Fig. 1b-d and 2a for additional information. **(d)** Graph of select mutants from (c). For each mutant, the log2 fraction of normalized reads, normalized to wild type, is plotted over time. The slope of a linear regression represents the selection coefficient. Shown is an average of n = 3 biological replicates. **(e-f)** Box and whisker plots showing the selection coefficients of groups of mutants with the indicated number of phosphomimetic mutations (e), or with specific substitutions at the indicated positions (f). The black center line indicates the median, boxes indicate the 25^th^-75^th^ percentiles, black lines represent 1.5 interquartile range (IQR) of the 25^th^ and 75^th^ percentile, black circles represent outliers. Data from n = 3 biological replicates are included. **(g)** Selection coefficients of indicated mutants. Error bars show standard deviation from the mean, shown is an average of n = 3 biological replicates. Significance was tested using repeated measures one-way ANOVA with Dunnett’s multiple comparisons test. Asterisks indicate significantly different from wild type (SC = 0), ****p < 0.0001, ***p < 0.0005, ns = non-significant.

To evaluate changes in the abundance of individual mutants, we calculated the log2 fold change in the fraction of reads, with respect to the initial population, for each mutant over time. Changes were then normalized to wild type and the data was visualized in a heat map (Fig. 2c). Importantly, the *hcm1-8A* mutant was depleted over time while the *hcm1-8E* mutant was enriched, validating our pooled approach. Clustering of the mutants based on the number of phosphomimetic mutations revealed several important features of phosphoregulation (Fig. 2c). First, fitness tended to increase with the number of phosphomimetic substitutions. However, when we examined a cluster of mutants that contained the same number of phosphomimetic sites (such as 4E, 4A) a wide range of fitness values was observed. This data rules out the possibility that a minimum number of phosphates is all that is needed for activation and argues that phosphorylation at particular sites can have a greater impact on activity and fitness. Importantly, when we considered the group of mutants that have only one phosphomimetic mutation (1E, 7A), we saw that no single site was sufficient to confer wild type fitness or better, suggesting that no single site is sufficient for activity. Conversely, all mutants that have only one unphosphorylatable mutation exhibited high fitness (7E, 1A), demonstrating that no single site is absolutely required for fitness (Fig. 2c; Extended Data Fig. 1c,d).

We next calculated a selection coefficient, or fitness score, for each mutant. The selection coefficient is the slope of the best fit line formed when the normalized log2 fraction of reads versus time is plotted for each mutant (Fig. 2d)^27^. Mutants that are less fit than wild type, like *hcm1-8A*, have a negative selection coefficient, whereas mutants that are more fit than wild type, like *hcm1-8E*, have a positive selection coefficient. Mutants with fitness similar to wild type have selection coefficients close to zero. To quantitatively examine how the number of phosphomimetic substitutions contributes to fitness, we calculated median selection coefficients for all mutants with a given number of phosphomimetic substitutions (Fig. 2e). These results confirmed that fitness increased with the number of phosphomimetic substitutions, with almost all mutants with six or more sites displaying increased fitness compared to wild type.

Next, we investigated the impact of phosphorylation at individual sites by calculating the median of the selection coefficients of all mutants with a specific substitution at each site (Fig. 2f). Overall, these data revealed that when all eight sites were mutated, glutamic acid substitutions increased fitness while alanine mutations reduced fitness at any site. However, mutations at two central sites, T460 and S471, had a disproportionally large impact on fitness.

In fact, phosphomimetic mutations at only sites T460 and S471 were sufficient to recapitulate wild type-like fitness (AAAEEAAA, Fig. 2d,g). Phosphomimetic substitutions at either of these sites individually, as the sole phosphomimetic site, resulted in an intermediate increase in fitness when compared to *hcm1-*8A (AAAEAAAA and AAAAEAAA, Fig. 2g). Conversely, blocking phosphorylation at either site individually reduced fitness when compared to *hcm1-*8E, with T460 having a larger effect (EEEAEEEE and EEEEAEEE, Fig. 2g). Interestingly, although T460 and S471 contributed the greatest amount to fitness, phosphorylation of these sites did not appear to be required, since a mutually exclusive phosphomutant that lacks phosphomimetic mutations at only these sites was also similar in fitness to wild type (EEEAAEEE, Fig. 2d,g). Together, these data indicate that both the overall number and the positions of phosphorylated sites can impact cellular fitness.

### Fitness levels predict the transcriptional activation and cell cycle regulatory functions of Hcm1

We next investigated whether the observed cellular fitness phenotypes correlated with Hcm1 activity. We previously found that the fitness phenotypes associated with *hcm1-8A* and *hcm1-8E* correlate with the decreased and increased expression, respectively, of Hcm1 target genes *in vivo*^25^. In addition, recruitment of Hcm1-8A to target gene promoters is reduced compared to wild type, although it is not eliminated. However, it remains unclear if TAD phosphorylation primarily regulates Hcm1 DNA binding, or if it stimulates its transcriptional activation function in another way. To separate these two possibilities and examine the effects of phosphosite mutations on Hcm1 activity, we utilized a transcriptional activation assay. The previously described Hcm1 TAD excludes the forkhead DNA-binding domain (DBD) but includes all eight C-terminal phosphosites of interest^28^. We fused wild type or phosphomutant TAD regions to the C-terminus of the LexA DBD and expressed these proteins in a reporter strain that contains eight LexA binding sites integrated upstream of the *lacZ* gene^29, 30^. β-galactosidase activity assays confirmed that the wild type Hcm1 TAD strongly activates transcription (Fig. 3a). Strikingly, expression of *hcm1-8A*, which cannot be phosphorylated, abolished all detectable activity. This demonstrates that phosphorylation can activate transcription independently of any effect on DNA binding. We next examined the activities of several phosphomutants from the screen, to determine if their activities correlated with fitness. Introduction of a single phosphomimetic mutation at either T460 or S471 (AAAEAAAA and AAAAEAAA) restored low levels of activity and phosphomimetic mutations at both key sites (AAAEEAAA) was sufficient to restore wild type-like activity. In addition, the mutant that only lacks phosphomimetic mutations at sites T460 and S471 (EEEAAEEE) displayed activity similar to wild type, consistent with its ability to restore near-wild type fitness (Fig. 2g). Overall, fitness values determined by Phosphosite Scanning were a good predictor of reporter gene activation. These data suggest that CDK phosphorylation of the TAD tunes Hcm1 activity by recruiting transcriptional activation machinery to stimulate target gene expression.

**Fig. 3.**
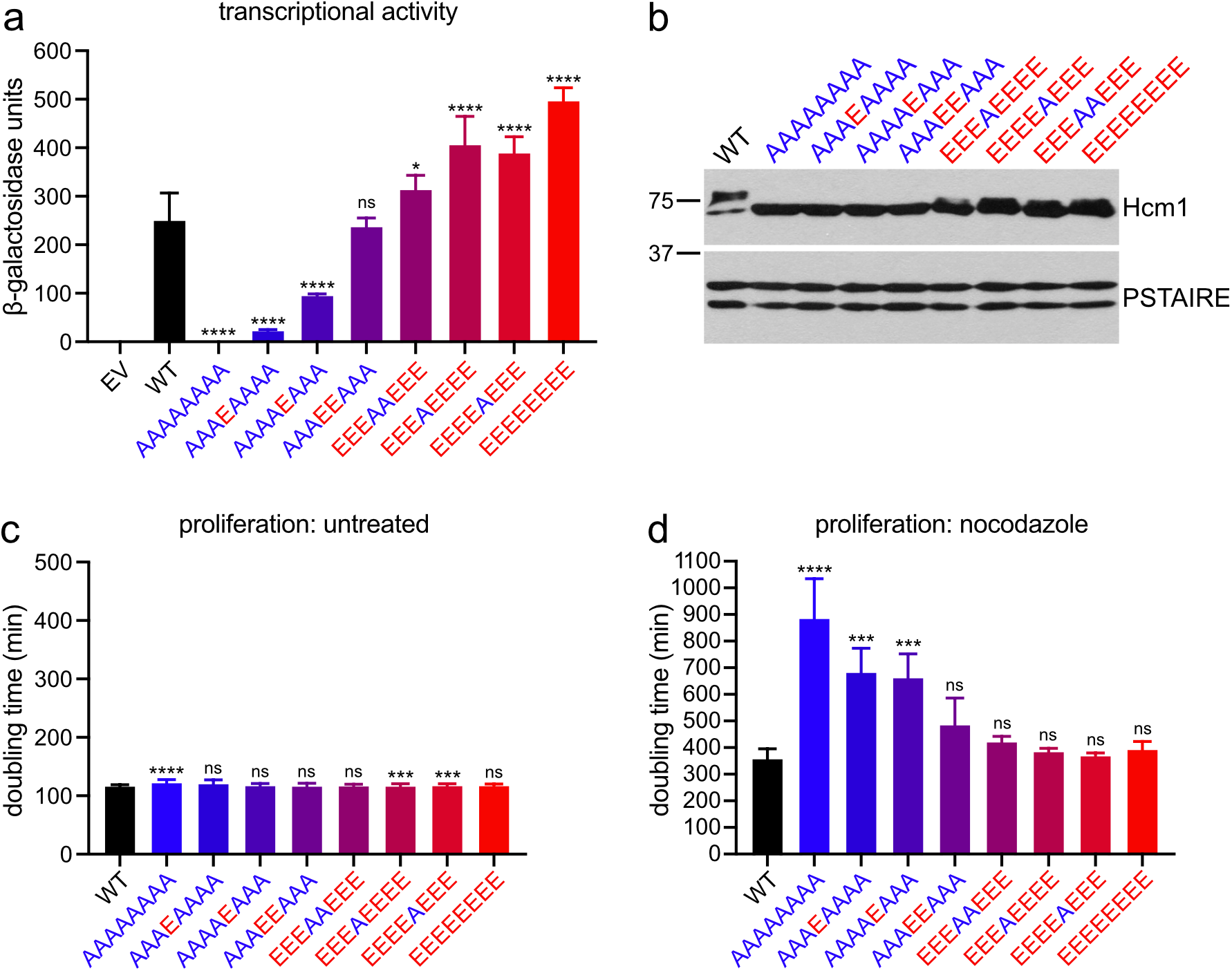
Fitness values predict the transcriptional activation and cell cycle regulatory functions of Hcm1. **(a)** Transcriptional activation assay of constructs containing the LexA-DBD fused to the Hcm1 TAD with the indicated phosphosite mutations, and wild type (WT) or empty vector (EV) controls. Error bars represent standard deviation, shown is an average of n = 3 replicates. Significance was tested using repeated measures one-way ANOVA with Dunnett’s multiple comparisons test. Asterisks indicate significantly different from wild type, ****p < 0.0001, *p < 0.05, ns = non-significant. See Extended Data Fig. 3a for additional information. **(b)** Western blot of the indicated Hcm1 proteins. Hcm1 was detected using an antibody against the C-terminal 3V5 tag, PSTAIRE shown as a loading control. **(c-d)** Doubling times of strains from (b) growing at 30°C in rich medium (c) or in rich medium with 5 μg/mL nocodazole (d). Error bars represent standard deviation, shown is an average of n = 3 replicates. Significance was tested using repeated measures one-way ANOVA with Dunnett’s multiple comparisons test. Asterisks indicate significantly different from wild type, ****p < 0.0001, ***p < 0.0005, ns = non-significant.

Next, we determined if the observed fitness phenotypes correlated with Hcm1 function *in vivo*. Hcm1 regulates several genes that are required for mitotic spindle function and as a result *hcm1Δ* cells are highly sensitive to microtubule poisons^24, 31, 32^. We therefore tested whether quantitative changes in Hcm1 activity imposed by different combinations of phosphosite mutations imparted similar changes in nocodazole sensitivity. Select phosphomutants were integrated into the genome at the *HCM1* locus and expressed at endogenous levels (Fig. 3b). Only modest differences in doubling times were observed among the phosphomutant strains grown in the absence of nocodazole (Fig. 3c). This result was expected since this measurement of proliferation is less sensitive than co-culture assays. However, when cells were grown in a sub-lethal concentration of nocodazole, we observed phosphorylation dependent sensitivity (Fig. 3d). Although nocodazole treatment slowed the growth of all cells, *hcm1-8A* cells grew approximately three-times slower than wild type in this condition. Mutants that mimic phosphorylation at either T460 or S471, but lack phosphorylation at all other sites (AAAEAAAA and AAAAEAAA), showed intermediate sensitivity, correlating with the increased fitness imparted by T460 or S471 phosphomimetic mutations. Moreover, mimicking phosphorylation at both T460 and S471 (AAAEEAAA), or at all sites *except* T460 and S471 (EEEAAEEE), was sufficient to restore nocodazole sensitivity to wild type levels. Together, these data demonstrate that the fitness phenotypes of Hcm1 unphosphorylatable and phosphomimetic mutants correlate with their cell cycle regulatory functions *in vivo*, and that both the position and number of phosphorylations contribute to Hcm1 activity.

### Stabilization of Hcm1 partially mitigates the effects of TAD phosphorylation

In addition to the TAD, CDK also phosphorylates a three-site phosphodegron in the Hcm1 N-terminus that triggers ubiquitin-mediated protein degradation^25^. Blocking phosphorylation of these sites with alanine substitutions (*hcm1-3N*) stabilizes Hcm1 and lengthens its expression through the cell cycle (Fig. S4A)^25^. We hypothesized that maintaining low levels of Hcm1 might be important for cells to respond to dynamic phosphorylation of the TAD, and as a result, phosphomimetic mutations might not enhance cellular fitness to the same extent in the *hcm1-3N* mutant, in which Hcm1 levels are stabilized throughout the cell cycle.

To test this hypothesis, we performed pairwise co-culture assays to evaluate the fitness effects of unphosphorylatable and phosphomimetic mutations in the TAD when Hcm1 is stabilized. First, we confirmed that *hcm1-3N* increases fitness compared to wild type, as previously reported^26^ (Fig. 4a). Next, we compared *hcm1-3N* to *hcm1-3N8A* and found that blocking phosphorylation of the TAD also reduced fitness when Hcm1 was stabilized (Fig. 4b). This result is consistent with the finding that *hcm1-8A* is unable to activate transcription (Fig. 3a). However, in contrast to the fitness benefit of *hcm1-8E* compared to wild type (Fig. 1e), *hcm1-3N8E* did not increase cellular fitness compared to *hcm1-3N* (Fig. 4c), in support of the hypothesis that TAD phosphorylation is less beneficial when Hcm1 is stabilized. These data argue that the balance between protein levels and activating phosphorylations is important for optimal regulation of Hcm1 activity.

**Fig. 4.**
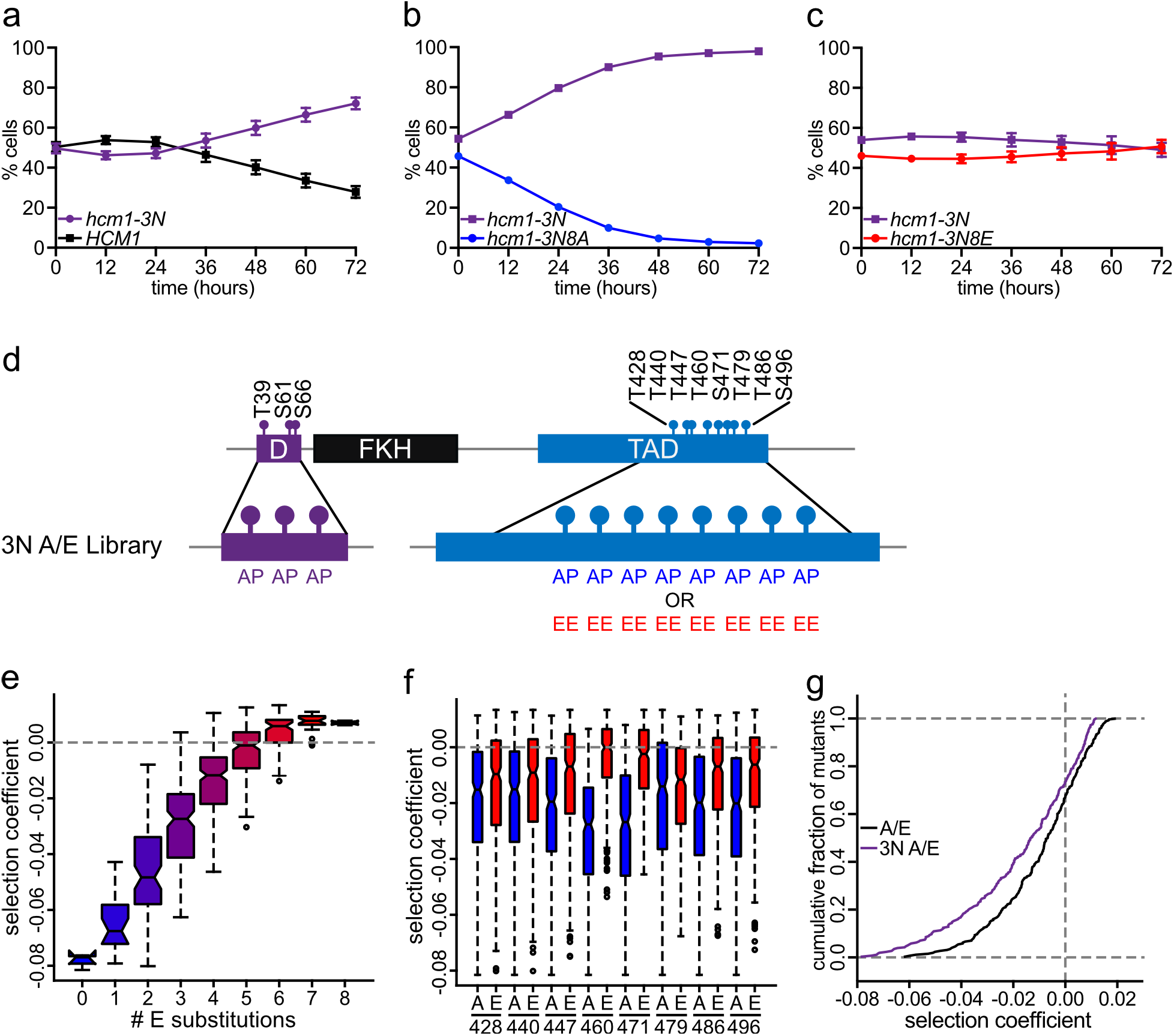
Stabilization of Hcm1 partially mitigates the effects of TAD phosphorylation. **(a-c)** Strains expressing the indicated *HCM1* alleles from plasmids were co-cultured and the percentage of cells expressing each allele was determined at the time points indicated. Shown is an average of n = 3 experiments, error bars represent standard deviations. **(d)** Schematic showing the design of the 3N A/E phosphomutant library. D indicates the phosphodegron consisting of three S/T-P sites that have been mutated to A-P (3N). The eight S/T-P sites in the TAD have been mutated to either A-P or E-E in all 256 possible combinations. **(e-f)** Box and whisker plots showing the median selection coefficients of groups of mutants with the indicated number of phosphomimetic mutations (e), or with specific substitutions at the indicated positions (f). The black center line indicates the median, boxes indicate the 25^th^-75^th^ percentiles, black lines represent 1.5 IQR of the 25^th^ and 75^th^ percentile, black circles represent outliers. Shown is an average of n = 3 biological replicates. See Extended Data Fig. 2b and 4 for additional information. **(g)** Cumulative frequency plot showing the cumulative fraction of mutants that display selection coefficients that are less than or equal to the indicated values. Black represents the A/E library, purple represents the 3N A/E library. Note that selection coefficients in the A/E library are calculated with respect to wild type *HCM1* and selection coefficients in the 3N A/E library are calculated with respect to *hcm1-3N*. Shown is an average of n = 3 biological replicates.

To investigate this result more comprehensively, we performed a second Phosphosite Scanning screen utilizing a phosphomutant plasmid library in which the A/E mutations were introduced into an *HCM1* allele that contained stabilizing mutations in the N-terminus (referred to as the 3N A/E library, Fig. 4d). This screen was performed in triplicate and analyzed for cellular fitness outcomes. However, since all *HCM1* alleles that were screened included stabilizing mutations, the calculated selection coefficients were normalized to the *hcm1-3N* allele (not wild type *HCM1*). We found that in the presence of stabilizing mutations, fitness also increased with the number of phosphomimetic mutations in the TAD (Fig. 4e). Interestingly, in contrast to what we observed in pairwise experiments, several mutants with large numbers of phosphomimetic substitutions, including *hcm1-3N8E*, were slightly more fit than *hcm1-3N* when the library was screened in bulk (Extended Data Fig. 4). However, for all phosphomimetic mutants, increases in fitness in the 3N background were reduced relative to the wild type (non-3N) background. For example, the average selection coefficient of the 8E mutant was 0.007 in the 3N A/E screen and 0.016 in the A/E screen. This suggests that our sequencing-based screens may be more sensitive than pairwise competition assays.

We next compared the importance of individual phosphosites in the 3N background and found that, similar to the results from the A/E screen, mimicking phosphorylation at any site increased fitness, with sites T460 and S471 having the largest impact (Fig. 4f). Interestingly, when we compared the results of the A/E and 3N A/E screens, we found that the fitness of both phosphomimetic and phosphodeficient mutants were lower in the 3N background, such that the positive effects of most phosphomimetic mutations were partially blunted and the negative effects of most phosphodeficient mutations were exacerbated (Extended Data Fig. 4). As a result, a larger fraction of 3N A/E mutants had a negative effect on fitness compared to A/E mutants (Fig. 4g). This effect was strongest among the least fit mutants. Together, these data support the model that increasing the levels of Hcm1 reduces the importance of TAD phosphorylation for Hcm1 activity and fitness.

### Quantifying fitness of mutants that incorporate wild type phosphosites

After identifying phosphomimetic mutations that activate Hcm1, we wanted to investigate if these sites are phosphorylated *in vivo*. To accomplish this, we performed two additional Phosphosite Scanning screens to interrogate phosphomutant plasmid libraries that incorporate wild type sites. We reasoned that comparing the fitness outcomes of mutants that incorporate wild type sites and analogous mutants where the phosphorylation status is fixed (A/E library) would allow us to infer which sites are phosphorylated *in vivo*.

In the first screen, we constructed a library in which each phosphoacceptor site was mutated to alanine or left as the wild type S-P or T-P site, in all possible combinations (referred to as the WT/A library, Fig. 5a). If all sites contribute to Hcm1 activity *in vivo*, our expectation was that all phosphomutants in this library should display either wild type-like or deleterious fitness. Consistent with this prediction, examination of each mutant’s abundance over time confirmed that all mutants displayed cellular fitness that was equal to or worse than wild type (Extended Data Fig. 5a). In general, fitness increased with the total number of available wild type sites (Fig. 5b), and a wild type S/T-P site at any location increased fitness (Fig. 5c). These data suggest that all eight sites are phosphorylated *in vivo* and contribute to Hcm1 activation.

**Fig. 5.**
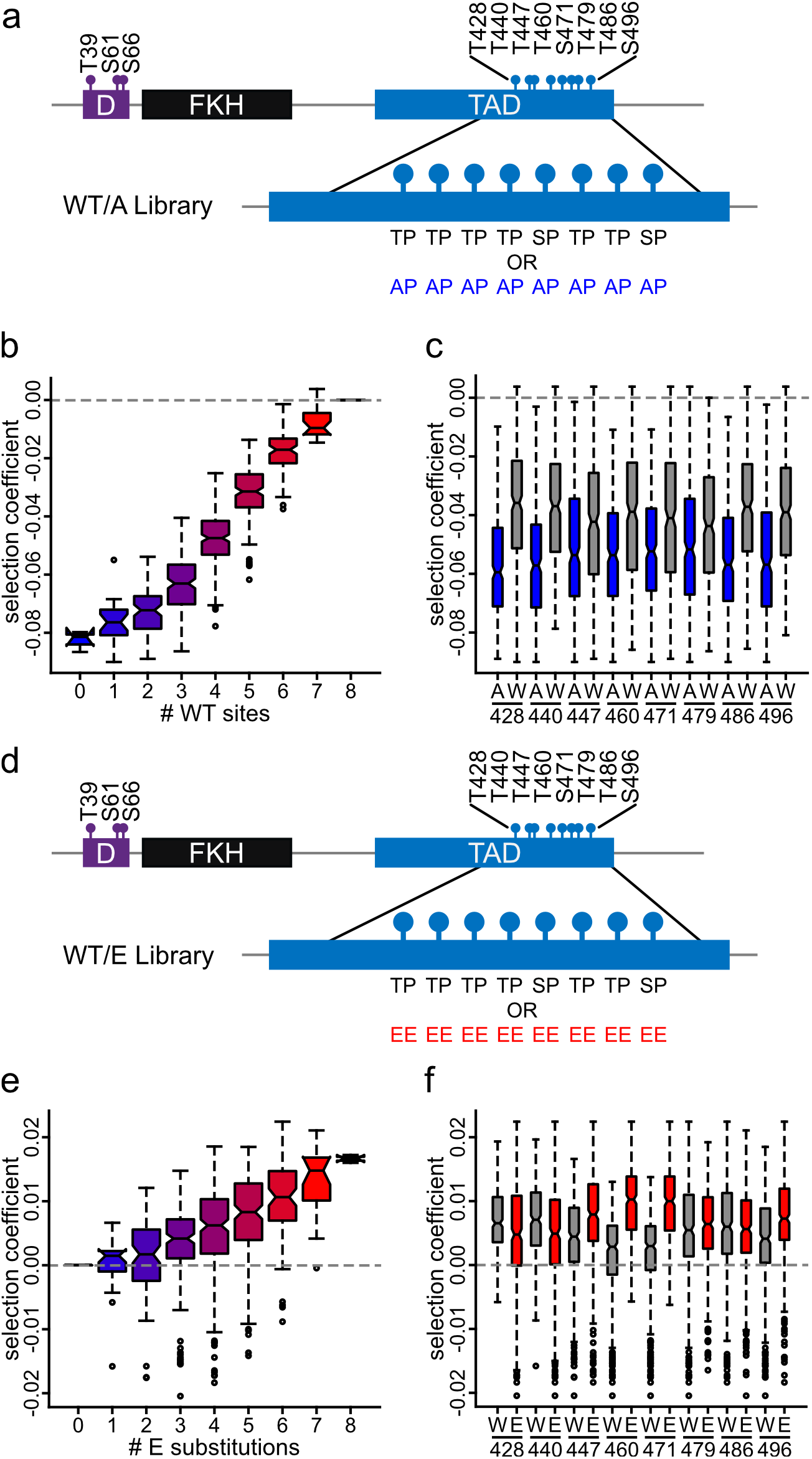
Quantifying fitness of mutants that incorporate wild type phosphosites. **(a)** Schematic showing the design of the WT/A plasmid library. S/T-P sites within the TAD were either left intact or mutated to A-P in all possible combinations (256). **(b-c)** Box and whisker plots showing the selection coefficients of groups of mutants with the indicated number of wild type (W) S/T-P sites (b), or with specific substitutions at the indicated positions (c). The black center line indicates the median, boxes indicate the 25^th^-75^th^ percentiles, black lines represent 1.5 IQR of the 25^th^ and 75^th^ percentile, black circles represent outliers. Shown is an average of n = 3 biological replicates. See Extended Data Fig. 2c and 5a for additional information. **(d)** Schematic showing the design of the WT/E plasmid library. S/T-P sites in the TAD were either left intact or mutated to E-E in all possible combinations (256). **(e-f)** Box and whisker plots showing the selection coefficients of groups of mutants with the indicated number of phosphomimetic mutations (e), or with specific substitutions at the indicated positions (f). The black center line indicates the median, boxes indicate the 25^th^-75^th^ percentiles, black lines represent 1.5 IQR of the 25^th^ and 75^th^ percentile, black circles represent outliers. Shown is an average of n = 3 biological replicates. See Extended Data Fig. 2d and 5b,c for additional information.

In the second screen, we constructed a library in which each site was either wild type or a phosphomimetic mutation (referred to as the WT/E library, Fig. 5d). Based on our previous observations, we expected to see a fitness benefit in highly phosphomimetic mutants. Indeed, we found that selection coefficients tended to increase with the number of phosphomimetic sites, similar to what we observed in previous screens (Fig. 5e). In fact, the median selection coefficient of all groups of WT/E mutants with one or more phosphomimetic sites was higher than that of wild type.

Next, we wanted to investigate the impact of phosphomimetic substitutions in more detail. Because phosphomimetic mutations at any site were beneficial compared to alanine substitutions, and all phosphosites in the library are wild type or phosphomimetic, we expected that all individual phosphomutants in this library would exhibit fitness that is equal to or better than wild type. Remarkably, several mutants were much less fit than wild type (Extended Data Fig. 5b). Examination of these mutants suggested that fitness phenotypes might be driven by E substitutions at one or more of the first three phosphosites (Extended Data Fig. 5c). To investigate this possibility quantitatively, we compared median selection coefficients of mutants based on their genotype at each site. Indeed, in contrast to what we observed in A/E and 3N A/E screens, we found that mutation of the first two sites to glutamic acid reduced fitness (T428 and T440, Fig. 5f). These data suggest that phosphomimetic mutations at some phosphosites are detrimental for fitness when wild type sites are also present.

### Phosphorylation of sites 460 and 471 depends on upstream CDK sites

To investigate why phosphomimetic mutations at T428 and T440 led to a reduction in fitness, we compared the selection coefficients of several phosphomutants from the WT/E screen. Mutation of either T428 or T440 to glutamic acid resulted in a modest, but not significant, reduction in fitness with respect wild type (EWWWWWWW and WEWWWWWW, Fig. 6a). This effect was strengthened and significant when both sites were mutated simultaneously (EEWWWWWW). Similarly, mutation of the first three positions resulted in a loss of fitness of approximately the same magnitude (EEEWWWWW). These data raised the possibility that wild type sites C-terminal to these mutations, including the most impactful sites for Hcm1 activity, T460 and S471, may not be phosphorylated in these mutants. Interestingly, adding a fourth phosphomimetic mutation at position 460 restored fitness to wild type levels (EEEEWWWW), in support of the possibility that T460 is not phosphorylated if the three N-terminal sites within the TAD are changed to E. However, this mutant was less fit than *hcm1-8E*, suggesting that additional downstream sites are also not phosphorylated when upstream sites are mutated.

**Fig. 6.**
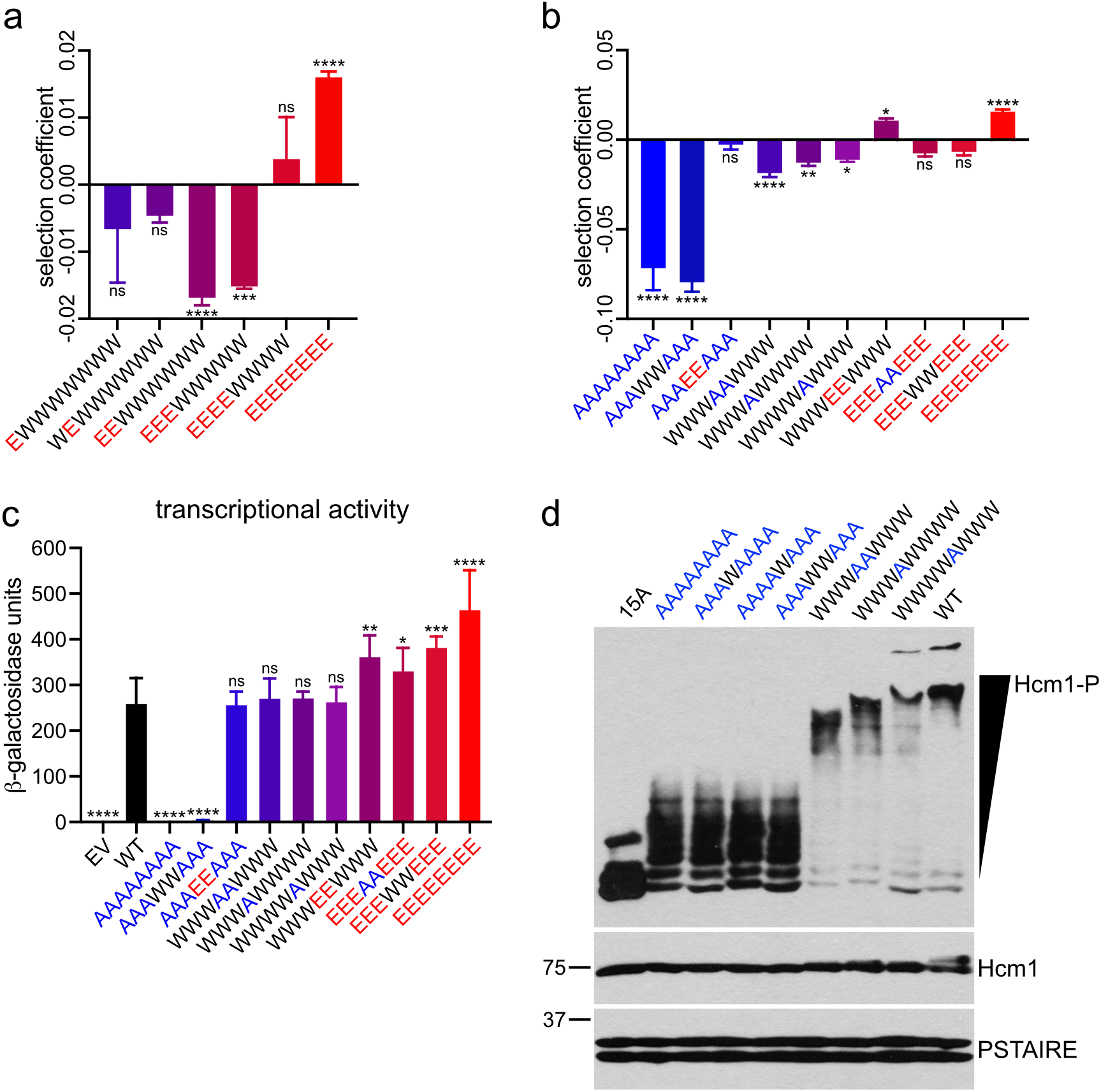
Phosphorylation of sites 460 and 471 depends on upstream CDK sites. **(a-b)** Selection coefficients of the indicated mutants from the WT/E screen (a) or multiple screens (b). Error bars represent standard deviation, shown is an average of n = 3 or 6 biological replicates. Significance was tested using ordinary one-way ANOVA with Dunnett’s multiple comparisons test. Asterisks indicate significantly different from wild type (SC = 0), ****p < 0.0001, ***p < 0.0005, **p < 0.005, *p <0.05, ns = non-significant. **(c)** Transcriptional activation assay of fusion constructs containing the LexA-DBD fused to the Hcm1 TAD with the indicated phosphosite mutations, or wild type (WT) or empty vector (EV) controls. Error bars represent standard deviations, shown is an average of n = 3 replicates. Significance was tested using repeated measures one-way ANOVA with Dunnett’s multiple comparisons test. Asterisks indicate significantly different from wild type, ****p < 0.0001, ***p < 0.0005, **p < 0.005, *p < 0.05, ns = non-significant. See Extended Data Fig. 3b for additional information. **(d)** Phos-tag Western blot of the indicated Hcm1 proteins (top panel) and standard Western blots showing expression of the indicated proteins (lower panels). Hcm1 was detected with antibody against the 3V5 tag, PSTAIRE is shown as a loading control.

One possibility is that although E substitutions mimic the charge of the phosphate and are sufficient for TAD activity, they may interfere with processive phosphorylation of more C-terminal sites because they are structurally dissimilar to a phosphate. If this is the case, then mutation of these sites to A would also be predicted to disrupt phosphorylation of more C-terminal sites. We examined several key mutants from the A/E, WT/A, and WT/E screens and found two comparisons that supported this model (Fig. 6b). First, we found in our initial screen that *hcm1-8A* has a severe fitness defect, and that mimicking phosphorylation at only sites T460 and S471 (AAAEEAAA) was sufficient to restore wild type fitness (Fig. 2g, 6b). We reasoned that if sites T460 and S471 are phosphorylated *in vivo* then a similar mutant with wild type sequence at only T460 and S471 (AAAWWAAA) should also show wild type-like fitness. However, this mutant showed a fitness defect approximately equal to *hcm1-8A*, suggesting the sites T460 and S471 are not phosphorylated when all other phosphosites are mutated to alanine (Fig. 6b.; compare three leftmost bars). These results were supported by transcriptional activation assays, which demonstrated that the AAAWWAAA mutant is unable to activate transcription (Fig. 6c). Second, our initial screen revealed that the fully phosphomimetic mutant (EEEEEEEE, *hcm1-8E*) displayed the largest fitness increase. Again, if T460 and S471 are phosphorylated *in vivo*, we would expect that a mutant with wild type sequence at these positions (EEEWWEEE) would display similar fitness and activity. However, we found that this mutant exhibited reduced fitness compared to *hcm1-8E* and was identical to the corresponding alanine mutant (EEEAAEE) (Fig. 6b; compare three rightmost bars). These two mutants (EEEWWEEE and EEEAAEEE) also showed similar transcriptional activity, which was reduced compared to *hcm1-8E* (Fig. 6c). Together, these results suggest that mutation of the first three phosphosites to either A or E impairs phosphorylation of T460 and S471, which in turn reduces Hcm1 activation.

Next, we sought to determine whether T460 and S471 are phosphorylated in the context of the wild type protein *in vivo*. Phosphomutants that only lack phosphoacceptor sites at T460 and/or S471 displayed a modest reduction in fitness, in support of this possibility (WWWAAWWW, WWWAWWWW, WWWWAWWW, Fig. 6b). To investigate phosphorylation more directly, we examined the migration of Hcm1 mutants using Phos-tag gels—which increase the mobility shift of proteins during SDS-PAGE in proportion to their phosphorylation levels—followed by Western blotting^33^. As previously shown^26^, most of the wild type Hcm1 protein migrates as a single band near the top of a Phos-tag gel (Fig. 6d). In contrast, a mutant that lacks all CDK consensus sites (15A) migrates as three bands near the bottom of the gel, confirming that the wild type protein is highly phosphorylated on S/T-P sites. Importantly, a mutant with alanine substitutions at T460 and S471 (WWWAAWWW) migrates faster than the wild type protein, and the corresponding single mutants are intermediate, providing strong evidence that T460 and S471 are phosphorylated *in vivo* (Fig. 6d; rightmost four lanes). Notably, there was no evidence of phosphorylation at T460 and/or S471 when all other sites were mutated to alanine. Mutants that are wild type at only T460, S471, or both positions migrate identically to the Hcm1-8A mutant that lacks all phosphosites in the TAD, supporting the possibility that mutation of the first three positions prevents phosphorylation at more C-terminal sites (Fig. 6d; leftmost five lanes). Together, these data demonstrate that T460 and S471 are phosphorylated *in vivo* and suggest that their phosphorylation depends upon the presence of wild type sequence at N-terminal CDK sites.

### Processive phosphorylation of sites 460 and 471 depends on Cks1

Since phosphorylation of T460 and S471 requires N-terminal CDK sites, we wondered if the phosphoadaptor protein Cks1 is involved in Hcm1 TAD phosphorylation. Cks1 forms a complex with cyclin-CDK and recruits the complex to already phosphorylated priming sites to trigger a processive multisite phosphorylation cascade in the N-to C-terminal direction^34–37^. Regulation by Cks1 requires a phosphorylated threonine priming site that is 12-30 residues upstream of additional CDK consensus sites. Interestingly, the spacing of CDK phosphosites in the Hcm1 TAD makes it a good candidate to be regulated by Cks1 (Fig. 7a). In addition, Cks1 appeared to facilitate processive Hcm1 phosphorylation in an *in vitro* kinase assay^36^. However, it is unknown if and how Cks1 regulates Hcm1 phosphorylation *in vivo*. Based on our phosphomutant analysis, we hypothesized that one or more of the first three CDK sites in the Hcm1 TAD acts as a priming site for Cks1, which in turn promotes downstream phosphorylation that contributes to Hcm1 activation.

**Fig. 7.**
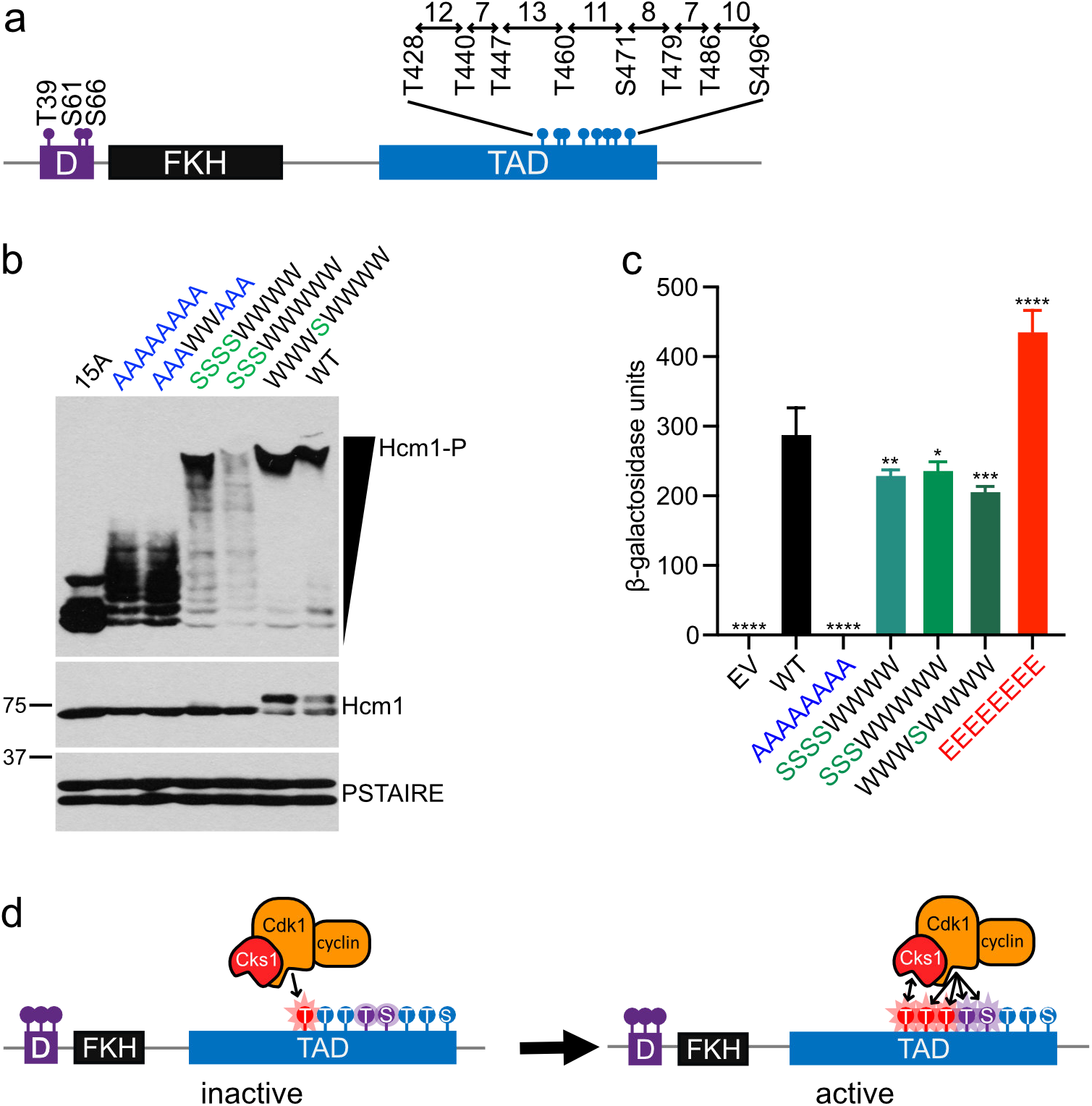
Processive phosphorylation of sites 460 and 471 depends on Cks1. **(a)** Schematic showing the spacing between phosphosites in the TAD. **(b)** Phos-tag Western blot showing phosphorylation of the indicated Hcm1 proteins (top panel) and standard Western blots showing expression of the indicated proteins (lower panels). Hcm1 was detected with antibody against the 3V5 tag, PSTAIRE is shown as a loading control. **(c)** Transcriptional activation assay of constructs containing the LexA-DBD fused to the Hcm1 TAD with the indicated phosphosite mutations or wild type (WT) or empty vector (EV) controls. Error bars represent standard deviation, shown is an average of n = 3 replicates. Significance was tested using repeated measures one-way ANOVA with Dunnett’s multiple comparisons test. Asterisks indicate significantly different from wild type, ****p < 0.0001, ***p < 0.0005, **p < 0.005, *p < 0.05. See Extended Data Fig. 3c for additional information. **(d)** Model depicting the role of Cks1 in multisite phosphorylation of the Hcm1 TAD.

Since Cks1 binds to phosphorylated threonines, but not phosphorylated serines^36, 37^, we examined Cks1 involvement in Hcm1 phosphorylation by changing T-P motifs to S-P motifs within the Hcm1 TAD. These T to S mutants can still be phosphorylated by CDK, but cannot serve as priming sites for Cks1. Strikingly, mutation of the first three or four threonine residues to serine reduced the processivity of Hcm1 phosphorylation, as evident by a ladder of intermediate migrating bands that were not observed with the wild type protein (SSSSWWWW and SSSWWWWW, Fig. 7b). Next, we tested if disruption of processive TAD phosphorylation resulted in a corresponding reduction in transcriptional activation by Hcm1. A modest, but significant, reduction in activity was observed in both mutants compared to wild type (Fig. 7c). We also considered the possibility that T460 could serve as a priming site to facilitate phosphorylation of S471 and/or downstream sites. There was no evidence of a reduction in processive phosphorylation when T460 alone was changed to serine (WWWSWWWW, Fig. 7b), however the activity of this mutant was reduced, suggesting that it may impact phosphorylation of one more C-terminal sites despite having a minimal impact on protein migration in a Phos-tag gel. Together, these results are consistent with a model in which the N-terminal threonine residues within the TAD can serve as Cks1 priming sites to facilitate the processive phosphorylation of downstream sites and enhance Hcm1 activity. These data further reveal that phosphomimetic mutations can substitute for bona fide phosphorylation for TAD activation, but not Cks1 recruitment.

## Discussion

### Processive phosphorylation by CDK activates Hcm1

Here, we describe a high-throughput selection approach, Phosphosite Scanning, that can be used to determine the contributions of individual phosphosites within a multisite phosphorylated domain. We used this approach to dissect the rules of phosphorylation within a model CDK-regulated domain, the TAD of the transcriptional activator Hcm1. First, by screening a library with all possible combinations of unphosphorylatable and phosphomimetic mutations, we were able to elucidate which combinations of phosphorylations activate Hcm1. This analysis revealed that Hcm1 activity is tunable and increases with the number of phosphomimetic mutations (Fig. 2e). However, not all sites are equivalent and some contribute a greater amount to activation (Fig. 2f). Remarkably, wild type-like Hcm1 activity can be achieved by non-overlapping patterns of phosphomimetic mutations (Fig. 2g, 3a, 3d). Although this screen revealed that it is possible to achieve activity in many ways, it did not elucidate which sites contribute to activation in the context of the wild type protein. For this reason, we also performed screens that incorporated wild type sites. Screening of the WT/A library revealed that mutation of any site to alanine reduces fitness, suggesting that all sites are phosphorylated to some extent *in vivo* (Fig. 5c). Screening of the WT/E library revealed more complex regulation. Phosphomimetic mutation of sites at the N-terminal end of the cluster prevented phosphorylation of more C-terminal sites (Fig. 5f). Our results suggest that the most N-terminal sites within the TAD both contribute to transcriptional activation and perform a structural role as docking sites for the CDK-associated processivity factor Cks1.

Together, these screens suggest a model for how the Hcm1 TAD is regulated by CDK *in vivo* (Fig. 7d). Our data suggest that CDK first phosphorylates an upstream threonine consensus site within the TAD region. This phosphorylated threonine then serves as a docking site for Cks1. Cks1 binds the phosphorylated priming site, triggering a multisite phosphorylation cascade in the N-to C-terminal direction. This allows for the rapid phosphorylation of two key sites, T460 and S471, which have the greatest impact on Hcm1 transcriptional activity, resulting in the expression of Hcm1 target genes. Notably, since the fully phosphomimetic mutant, *hcm1-8E*, displays greater activity and fitness than wild type Hcm1 (Fig. 2g, 3a), this suggests that, on average, fewer than eight sites are phosphorylated at any one time. Instead, it’s likely that the phosphorylation of the TAD is dynamic, since all sites contribute to activity to some extent (Fig. 5c).

Although Cks1 has been shown to enhance the phosphorylation of Hcm1 *in vitro*^36^, our results provide the first evidence that Hcm1 is processively phosphorylated by Cks1 *in vivo*. To facilitate processive phosphorylation, Cks1 binds to a phosphorylated threonine that is 12 to 30 amino acids upstream of additional CDK sites. Based on their spacing, any of the first four CDK sites in the Hcm1 TAD could act as priming sites for one or more downstream sites (Fig. 7a). In fact, all threonine residues in the TAD, with the exception of T440, were previously identified as putative Cks1-binding sequences^37^. Our expectation was that Cks1 binding at the N-terminal end of the cluster would have a greater impact on phosphorylation, since it could initiate a processive cascade of phosphorylation that would lead to rapid phosphorylation of T460 and S471 and activation of the protein. In support of this model, the extent of Hcm1 phosphorylation was reduced when the three N-terminal threonines within the TAD were mutated to serine, whereas the T460S mutation alone had less of an effect (Fig. 7b). Interestingly, although the T460S mutation alone did not impair Hcm1 phosphorylation as visualized by Phos-tag Western blot, the activity of this mutant was significantly reduced compared to wild type (Fig. 7b, c). Therefore, it is possible that T460S does impair phosphorylation of one or more downstream sites, but T460 phosphorylation may impact fewer sites than T428/T440/T447.

Phosphorylation of the Hcm1 TAD activates expression of Hcm1 target genes *in vivo*. Our data indicates that TAD phosphorylation stimulates transcriptional activation through a charge-based mechanism that is independent of DNA binding (Fig. 3a). A likely possibility is that phosphorylation causes a conformational change that promotes binding to a coactivator. Interaction with a coactivator could stabilize the Hcm1-DNA interaction, and this might explain why the unphosphorylatable *hcm1-8A* protein exhibits decreased recruitment to target gene promoters *in vivo*^25^. Our results suggest that a balance between the overall number and precise positioning of phosphorylated residues within the TAD determines cofactor binding.

Interestingly, the key sites T460 and S471 are centrally located within the phosphosite cluster and are in a region that is predicted to be more structured than the surrounding sequence (Fig. 1a). Analysis of yeast and human TADs have shown that negative charge is important in TAD to keep critical aromatic resides solvent exposed, which enables binding to coactivator^38, 39^. Therefore, phosphorylation of T460 and S471 may increase the negative charge in this critical region, leading to coactivator recruitment. This possibility is also consistent with the fact that a mutant that has phosphomimetic mutations at all sites except T460 and S471 (EEEAAEEE) can also support transcriptional activity, since an increased amount of negative charge that is less optimally placed could be sufficient for cofactor recruitment. We anticipate that this mechanism of regulation may be common among multisite phosphorylated substrates: that regulation can occur via different combinations of phosphosites, but that evolution has selected for sequence contexts that promote highly processive phosphorylation and switch-like regulation.

### Decoding multisite phosphorylation by Phosphosite Scanning

Historically, determining which phosphosites within a multiphosphorylated domain impact protein function has been challenging, since it requires making large numbers of mutant alleles and screening them individually. Phosphosite Scanning offers several benefits over traditional approaches. Since all possible combinations of alleles can be generated and screened in bulk, it is both comprehensive and rapid. Moreover, as is evident from our data, fitness readouts are very sensitive and can detect small differences in activity among mutants. The ideal substrate for this type of screen is a protein like Hcm1, in which eliminating and mimicking phosphorylation lead to opposing effects on protein function and cellular fitness. This approach can easily be applied to any phosphorylated protein that is regulated by multisite phosphorylation and demonstrates phosphorylation-dependent fitness effects. Phosphosite Scanning could also be adapted for substrates that do not meet these criteria. Instead of measuring cellular fitness as a readout, protein function could be measured in another way. For example, screens could be developed that utilize GFP as a readout, and mutants with different effects on activity determined by sorting cells based on GFP expression. This type of approach has been successful in combination with traditional deep mutational scanning^38, 39^.

One consideration when designing a Phosphosite Scanning screen is that the functional consequence of phosphorylation is not always replicated by phosphomimetic mutations. Substitution of phosphoacceptor sites with two glutamic acids replicates the change of charge caused by phosphorylation. This can mimic the outcome of phosphorylation when the change of charge of the substrate region is important for protein function^18^. However, there are instances when the addition of a phosphate group is required, and the effect cannot be replicated by charge alone. This is often the case when phosphorylation regulates a protein-protein interaction. For example, the degradation of many substrates of SCF ubiquitin ligases depends on substrate phosphorylation to form a phosphodegron motif that can interact with an F-box protein^16, 17^. Since glutamic acid substitutions are structurally dissimilar to a phosphate, they are unable to form an interaction motif. Similarly, interaction between a CDK substrate and the processivity factor Cks1 depends on a phosphorylated threonine priming site within the substrate^36, 37^. As our data shows, phosphomimetic substitutions at the most N-terminal threonine residues in the Hcm1 TAD prevented phosphorylation of more C-terminal wild type sites, suggesting that they do not interact with Cks1. Despite these limitations, screens of phosphomimetic mutations can be informative for many targets. Alternatively, variations on Phosphosite Scanning that only interrogate unphosphorylatable substitutions could also be used to determine which sites are required for a functional output.

### The importance of Hcm1 phosphoregulation

Hcm1 is an ideal model substrate to elucidate the principles of regulation by CDK because the TAD shares many features of domains that are multisite phosphorylated^3^, and its phosphorylation leads to clear fitness phenotypes that correlate extremely well with protein function. However, the fact that cells expressing the fully phosphomimetic mutant (*hcm1-8E*) are more fit than wild type cells raises the interesting question of why Hcm1 phosphoregulation has evolved. A likely possibility is that in some conditions having too much Hcm1 activity is detrimental, so it is beneficial to keep its levels low and rapidly toggle activity on and off via phosphorylation, as needed. In support of this model, activating phosphates in the TAD are specifically removed by the phosphatase calcineurin in response to stress^26^. We show here that phosphorylation-mediated degradation is important to sensitize Hcm1 to changes in TAD phosphorylation, since phosphomimetic mutations in the TAD do not provide the same fitness advantage when N-terminal phosphosites are mutated and the protein is stabilized (Fig. 4). The complex phosphoregulation of Hcm1 activity suggests that it is a critical regulator that modulates fitness in different environments. Future studies examining the consequence of Hcm1 phosphomutants in different environments should shed light on this possibility.

## Acknowledgements

The authors thank Peter Pryciak, Matthew Winters, Fritz Roth, Julia Flynn, Daniel Bolon, Sneha Gopalan and Tong Wu for reagents and technical advice. We also thank members of the Benanti lab for helpful discussions. This work was supported by National Institutes of Health grants R01GM117152 and R35GM136280 to J.A.B. and R01HD072122 to T.G.F.

## Author Contributions

M.M.C and J.A.B designed the study. M.M.C. and M.A.N.R. performed the experiments. R.L. generated the count tables from raw sequencing data. L.J.Z. supervised R.L. M.M.C., J.A.B. and T.G.F performed all additional analyses. M.M.C. and J.A.B. wrote the manuscript with input from all authors.

## Methods

### Yeast strains and plasmids

All strains are described in Supplementary Table 1. Genetic manipulations were performed using standard methods^40, 41^. Strains were grown in rich medium (YM-1) or synthetic medium lacking a single amino acid with 2% dextrose or galactose, where indicated, at 30°C. Strains with *HCM1* mutations integrated into the genome were generated from synthesized gene fragments (Invitrogen) that were cloned into pFA6a-3V5-KanMX6 using the NEBuilder HiFi DNA Assembly kit (New England Biolabs). Strains for pairwise competition assays were generated by inserting a *Hyg-TEFp-GFP* expression cassette into a gene-free region of chromosome VI^42^. A previously characterized non-fluorescent GFP mutant (GFP-Y66F) was integrated in place of wild type GFP and used as a matched negative control^43^.

All plasmids are listed in Supplementary Table 2. To generate *HCM1* expression plasmids, the *HCM1* open reading frame and endogenous promoter were amplified from the genome and cloned into pRS316. Plasmids for β-galactosidase transcriptional activation assays were generated using the pBTM116 backbone. The mutant or wild type *HCM1* TAD region (amino acids 306-511) was amplified by PCR from genomic DNA. The C-terminal 3V5 tag was PCR amplified from the pFA6a-3V5-KanMX6 vector. Both products were purified and cloned pBTM116 using the NEBuilder HiFi DNA assembly kit (New England Biolabs).

Phosphomutant libraries used for Phosphosite Scanning screens were cloned into a pRS316 vector containing the endogenous *HCM1* promoter upstream of the *HCM1* open reading frame (wild type or *hcm1-3N*) in which the region to be mutated had been deleted and replaced by an SphI restriction enzyme site (pRS316-HCM1*SphI or pRS316-hcm1-3N*SphI), to facilitate HiFi DNA assembly.

### Pairwise co-culture competition assays

For pairwise co-culture assays, *GAL1p-HCM1* strains expressing wild type GFP or non-fluorescent GFP-Y66F were transformed with pRS316 plasmids expressing the indicated *HCM1* alleles and cultured in synthetic media lacking uracil with 2% galactose. Logarithmic phase cells were mixed in equal proportions by diluting 1 optical density of each strain into 10 mL media with galactose. 0.15 optical densities of the mixed culture were then collected, pelleted by centrifugation, resuspended in 2 mL sodium citrate buffer (50mM sodium citrate, 0.02% NaN3, pH 7.4) and stored at 4°C, for analysis by flow cytometry at the conclusion of the experiment.

Mixed cultures were diluted to 0.006 optical densities in synthetic media lacking uracil with 2% dextrose and allowed to grow for 12 hours, during which time the cultures did not reach saturation. Cultures were diluted in this manner every 12 hours for a total of 72 hours. At each timepoint, 0.15 optical densities were collected and resuspended in 2 ml sodium citrate buffer and stored at 4°C for subsequent analysis. At the conclusion of the time course, the percentage of each strain in each sample were determined by quantifying the percentage of GFP positive cells with a Guava EasyCyte HT flow cytometer. Data was analyzed with FlowJo software. N = 3 technical replicates were performed in each time course and percentages at each time point averaged together. Data presented is an average of n = 3 biological replicates.

### Western blotting

Lysates were prepared using TCA extraction and Western blotting and Phos-tag Western blotting was carried out as previously described in^44^. Proteins were visualized using antibodies that recognize PSTAIRE (P7962, Sigma), V5 (Invitrogen), and HA (12CA5).

### Phosphosite Scanning library construction

Phosphomutant plasmids were generated by annealing and extending overlapping mutagenic oligonucleotides spanning a 270 base pair region that encompasses all phosphosites, as detailed in Extended Data Fig. 1a. Six oligonucleotides were designed that cover one to three phosphosites each, depending on the distance between phosphosites, with 21-27 base pairs of overlap between consecutive oligonucleotides. For each of the oligonucleotides, several variations were designed to include wild type, alanine and glutamic acid substitutions at each phosphosite, in all possible combinations. See Supplementary Table 3 for oligonucleotide sequences. Equimolar ratios of the desired mutagenic oligonucleotides were then combined to a final concentration to 100 µM. For example, to generate the AE library, oligonucleotides encoding A and EE substitutions at each site were combined. 6 µM of pooled oligonucleotides were then phosphorylated at the 5’ ends by incubation with 1 mM ATP, 10 units T4 polynucleotide kinase (New England Biolabs), 1X T4 Polynucleotide Kinase Buffer (New England Biolabs) in a total volume of 50 µL at 37°C for one hour. Next, oligonucleotides were annealed by diluting 10 µL phosphorylated oligonucleotides to a final volume of 20 µL with sterile water, heating to 95°C and allowing to slowly cool to 25°C over the course of one hour. Single stranded gaps between hybridized oligonucleotides were then filled in by combining 5 µL of the hybridized oligonucleotides from the previous annealing step with 1 unit Q5 High Fidelity DNA polymerase (New England Biolabs), 1X Q5 Reaction Buffer (New England Biolabs), 0.5 mM dNTPs (New England Biolabs), and sterile water to a final volume of 10 µL and incubating at 72°C for 2 minutes. Gaps were then ligated by adding 20 units Taq DNA Ligase (New England Biolabs), Taq DNA Ligase Buffer (New England Biolabs), and sterile water to the 10 µL reaction from the previous step to a total volume of 15 µL. The reaction was incubated at 45°C for 20 minutes. Finally, the DNA template was amplified by PCR using Phusion High-Fidelity DNA polymerase and Phusion GC Buffer (New England Biolabs). The PCR product was purified using the DNA Clean & Concentrator-25 kit (Zymo Research) and recombined into and SphI digested pRS316-HCM1*SphI using the NEBuilder HiFi DNA assembly kit (New England Biolabs). Plasmids were then transformed into NEB 10-beta Competent *E. coli* (High Efficiency) (New England Biolabs). All plasmid libraries were constructed in triplicate and mixed prior to transformation to maximize coverage of phosphomutants and avoid bias in oligo mixture or annealing steps.

### Phosphosite Scanning Screens

Plasmid libraries that were constructed in triplicate were mixed in equal proportion for transformation into cells. Plasmid libraries were transformed into a *GAL1p-HCM1* strain grown in YM-1 containing 2% galactose, to maintain expression of endogenous *HCM1*. For each screen, wild-type or Hcm1-3N expressing plasmids were added to each library as a control.

Transformed cells were grown overnight in liquid C-Ura with 2% galactose at 23°C. After approximately 16 hours an aliquot of cells was removed and plated on C-Ura to confirm sufficient transformation efficiency (10X library coverage). The remaining cells were pelleted and washed five times with 15 mL C-Ura with 2% galactose to remove any untransformed plasmid and resuspended in 50 mL C-Ura with 2% galactose and allowed to grow to log phase (approximately 48 hours) at 30°C. Cells were then diluted to an optical density of 0.008 in 75 mL C-Ura with 2% galactose and grown overnight at 30°C. At the start of the experiment, the population was sampled to determine the representation of each plasmid in the population before selection. 20 optical densities were collected, frozen on dry ice and stored at -80°C for subsequent recovery of plasmids and preparation of sequencing libraries. One optical density of cells was harvested for Western blotting to confirm levels of plasmid and endogenous Hcm1.

Cells were then diluted to an optical density of 0.008-0.016 into C-Ura with 2% dextrose to repress expression of endogenous *HCM1* and allowed to grow for 12 hours at 30°C to mid-logarithmic phase. The population was sampled as above and diluted into C-Ura with 2% dextrose every 12 hours for a total of 72 hours.

### Illumina sequencing library preparation

Plasmids were recovered from frozen cell pellets using the YeaStar Genomic DNA Kit (Zymo Research). Plasmid *hcm1* sequence was amplified by PCR for 21 cycles with vector specific primers using Phusion High-Fidelity DNA polymerase (New England Biolabs). Products were extracted from a 1% agarose gel using the QIAquick Gel Extraction Kit (Qiagen). TruSeq sequencing adapters were added to mutant fragments by PCR using *HCM1* specific primers fused to the TruSeq universal adapter or unique TruSeq indexed adapters. Products were extracted from a 1% agarose gel using the QIAquick Gel Extraction Kit (Qiagen). Libraries were pooled and paired-end 150 base pair sequencing reads obtained by sequencing on a HiSeq4000 platform (Novogene). Due to high sequence identity between samples all libraries were mixed with at least 50% DNA of heterogeneous sequence to increase diversity.

### Phosphosite Scanning data analysis

All sequencing data is available from the NCBI Sequence Read Archive (BioProject # PRJNA841829). *HCM1* alleles were counted for each paired-end (PE) sequencing fragment that had an exact sequence match in both reads, using a custom Python script (Supplementary File 1). Count tables for all screens are included in Supplementary Dataset 1. Average data from replicate screens are presented in heat maps (Fig. 2c and Extended Data Fig. 1c-d, 4b-c, 5), which show the log2 fraction of reads at the indicated time points. Primary data plotted in heat maps are included in Supplementary Dataset 2. Selection coefficients were then determined for each mutant by calculating the slope of the log2 fraction of reads versus time for each mutant and subtracting the slope of the log2 fraction of reads versus time for wild type (or *hcm1-3N* for the 3N A/E screen). Selection coefficients for all screens are included in Supplementary Dataset 3. Median selection coefficients for select groups of mutants (with specific numbers of phosphomimetic substitutions, or with specific substitutions at each position) were then calculated and plotted in box and whisker plots using *boxplot()* in R. In all cases, values from all three replicates of each screen are included in the medians. Bar graphs comparing selection coefficients of select mutants show average values from each of three replicate screens.

### Doubling time assays

Cells were grown to mid logarithmic phase, then diluted to 0.1 optical densities in YM-1 with or without 5 µg/mL nocodazole. Cells were transferred to 96 well plates in triplicate and grown at 30°C with shaking in a Tecan Infinite M Nano plate reader. Optical densities at 600 nm were measures every 20 minutes for a total of 33 hours. Doubling times were quantified by fitting data points between 0.2 and 0.5 optical densities to an exponential growth equation using GraphPad Prism software. Multiple isolates were tested for each genotype listed.

### β-galactosidase transcriptional activation assays

Cells with an integrated *lacZ* reporter and carrying plasmids expressing LexA-DBD-Hcm1 fusion proteins were grown in synthetic media lacking tryptophan. For each strain, the optical density at 600 nm was taken and 1 mL of cells were collected. Cells were pelleted and the media removed. Cells were then resuspended in 0.5 mL Z-buffer (60 mM Na2HPO4 heptahydrate, 40 mM NaH2PO4 monohydrate, 10 mM KCl, 1mM MgSO4 heptahydrate) and permeabilized through the addition 50 µL chloroform and 10 µL 0.4% SDS followed by vortexing. 300 µL Z-buffer containing 2.4 mg/mL *o*-nitrophenyl-β-d-galactopyranoside and 6 µL/mL β-mercaptoethanol was then added and the time recorded. Reactions were mixed, moved to 30°C and monitored for a colorimetric change for a maximum of 120 minutes. Reactions were stopped either when a yellow color change was observed or after 120 minutes. To stop reactions, 0.5 mL 1M Na2CO3 was added with vortexing and the time recorded. Completed reactions were stored on ice until all were finished. Chloroform was removed by centrifugation and the optical density of 1 mL of aqueous supernatant was measured at 405 nm. β-galactosidase units were calculated as (1000*OD405)/(OD600*1 mL*reaction time [min]). All reactions were carried out in triplicate and averaged together. Average values from biological replicates were then averaged together and plotted where indicated.

## Extended Data Figures

**Fig 1. Supporting data for A/E screen.**

**Fig. 2. Replicate correlation plots for all screens.**

**Fig. 3. LexA-Hcm1 fusion proteins are expressed at similar levels.**

**Fig. 4. Supporting data for 3N A/E screen.**

**Fig. 5. Supporting data for the WT/A and WT/E screens.**

## Supplementary Information

### Supplementary Tables

**Table 1. Strain table.**

**Table 2. Plasmid table.**

**Table 3. Oligonucleotide table.**

### Supplementary Files

**File 1. Python code used to generate count tables.**

### Supplementary Datasets

**Dataset 1. Count tables for all screens.**

**Dataset 2. Primary data for heatmaps.**

**Dataset 3. Selection coefficients for all screens.**

