## Supplementary Information for "Phosphosite Scanning reveals a complex phosphorylation code underlying CDK-dependent activation of Hcm1"

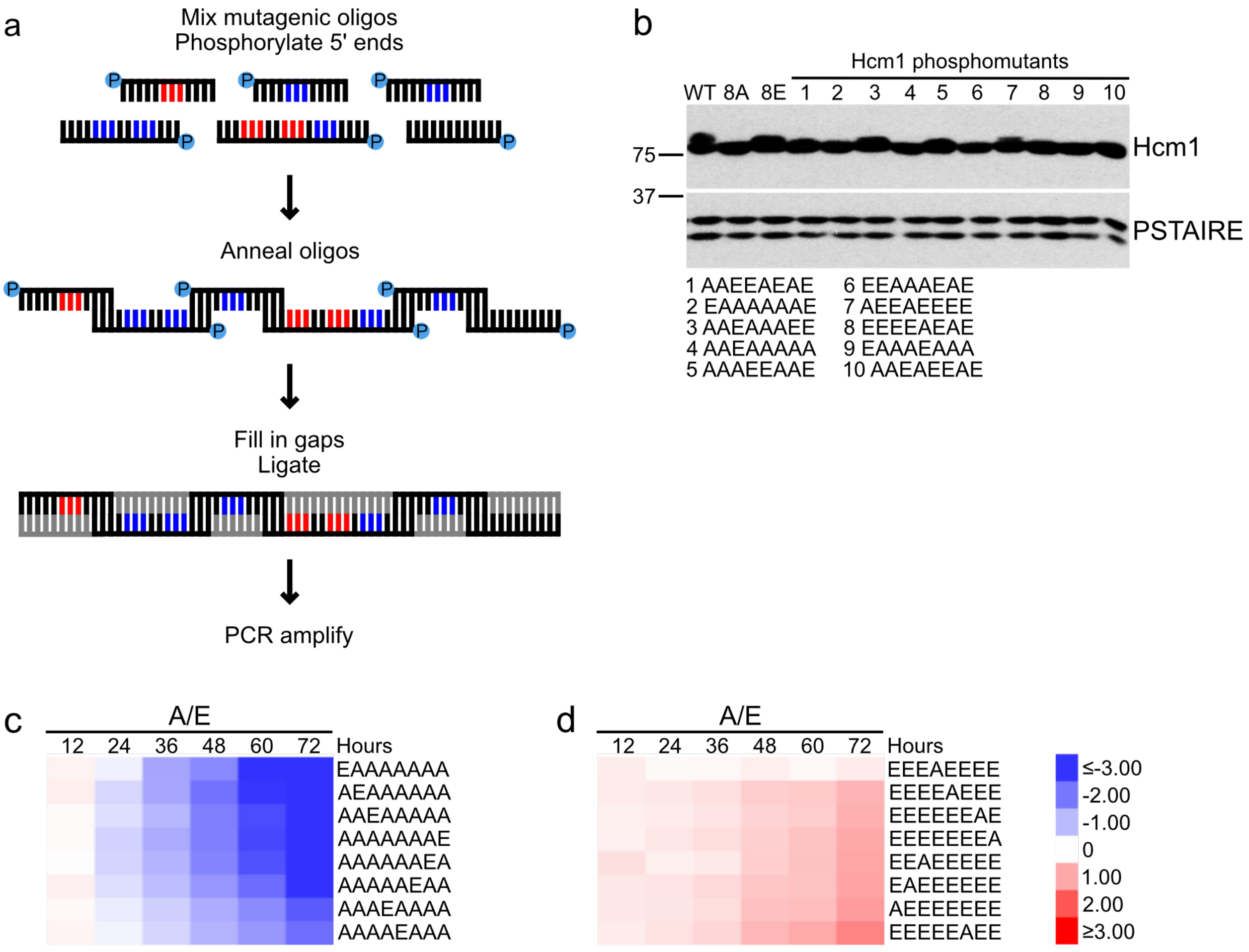


**Extended Data Fig. 1. Supporting data for the A/E screen. (a)** Schematic depicting the construction of phosphomutant plasmid libraries. See Methods for more details. **(b)** Western blot of representative Hcm1 mutants and wild type (WT) proteins from the A/E screen. Hcm1 was detected using antibody recognizing a C-terminal 3V5 tag, PSTAIRE is shown as a loading control. **(c-d)** Expanded view of 7A, 1E (c) and 1A, 7E (d) clusters from Fig. 2c. Each row represents a mutant with the indicated phosphomimetic mutations, shown is the log2 fold change in normalized read counts with respect to time zero for each mutant, all mutants have been normalized to wild type. Blue indicates depletion, red indicates enrichment. Shown is an average of n = 3 biological replicates. Scale bar is the same for (c) and (d).

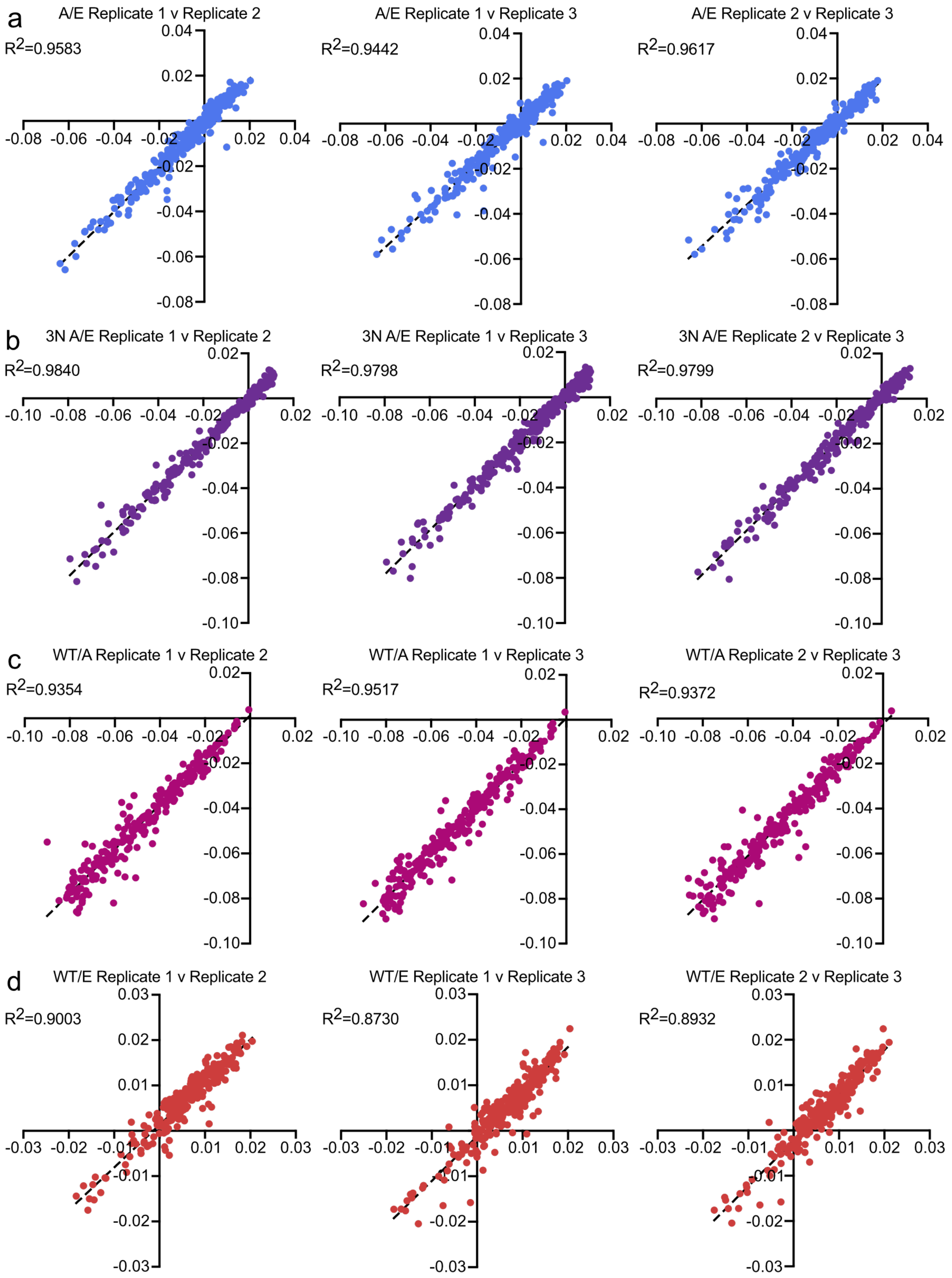


**Extended Data Fig. 2. Replicate correlation plots for all screens.** Scatter plots showing the correlation between independent replicates from the A/E **(a)**, 3N A/E **(b)**, WT/A **(c)**, WT/E **(d)** screens. $R^2$ values are shown with each plot.

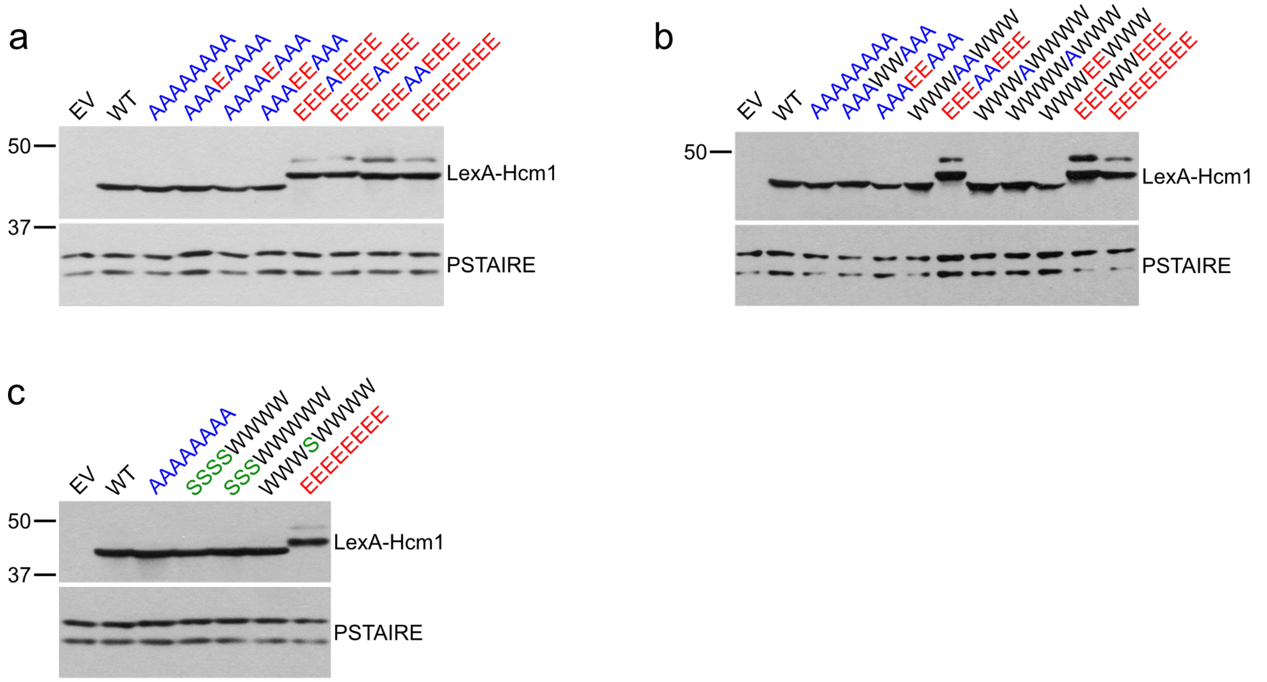


**Extended Data Fig3. LexA-Hcm1 fusion proteins are expressed at similar levels.** Western blot showing expression of LexA-Hcm1 fusion constructs of the indicated genotypes to support Fig. 3a **(a)**, 6c **(b)**, and 7c **(c)**. Wild type (WT) and empty vector (EV) controls are included. LexA-Hcm1 was detected using antibodies that detect a C-terminal 3V5 tag, PSTAIRE is shown as a loading control.

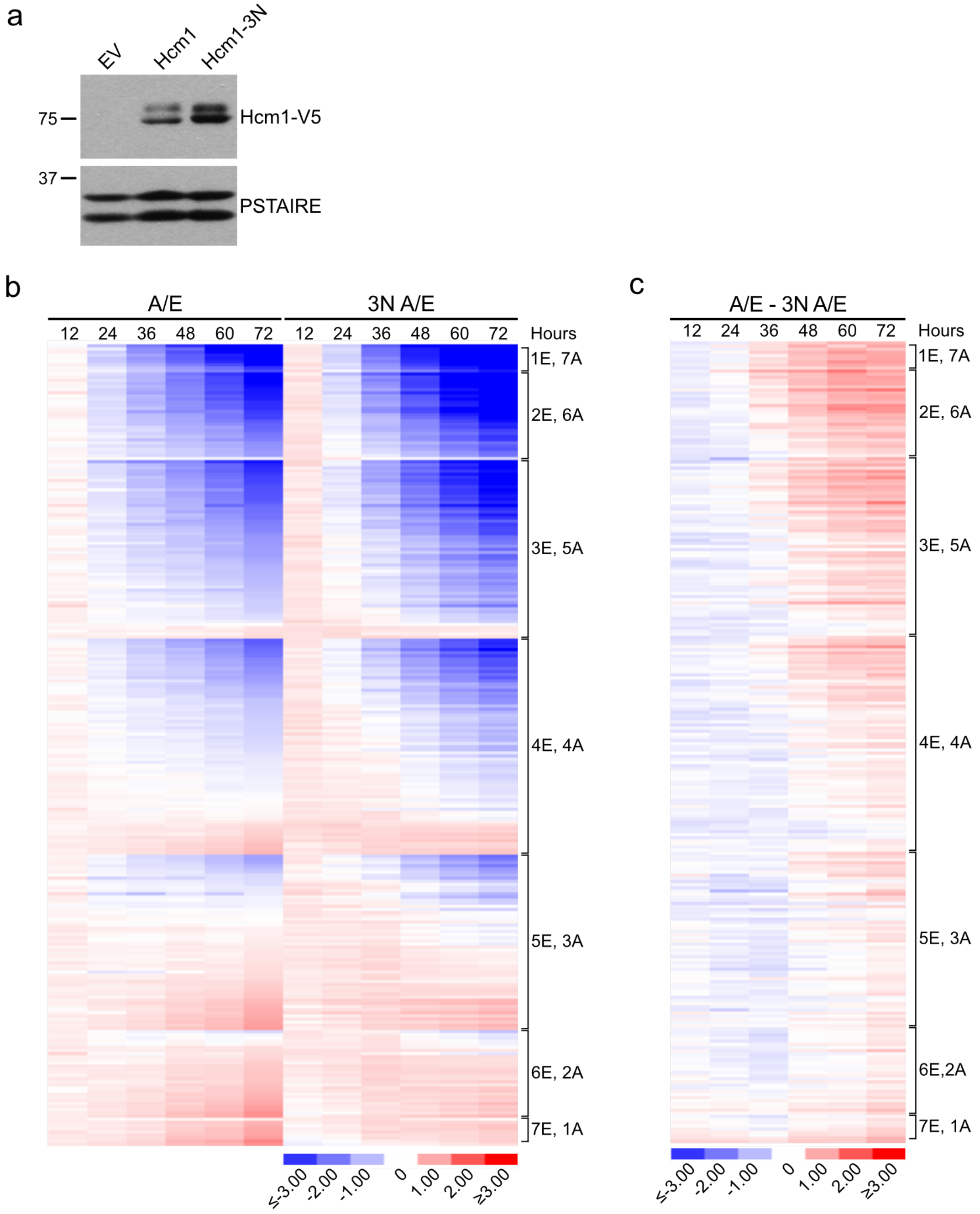


**Extended Data Fig. 4. Supporting data for the 3N A/E screen. (a)** Western blot showing expression of Hcm1 and Hcm1-3N from low copy plasmids (pRS316). Hcm1 was detected using antibodies that recognize a C-terminal 3V5 tag, PSTAIRE is shown as a loading control. **(b)** Side-by-side heat map representation of the results of the A/E and 3N A/E Phosphosite Scanning screens. Each row represents a mutant, shown is the log2 fold change in normalized read counts with respect to time zero for each mutant. In the A/E screen all mutants were normalized to WT. In the 3N A/E screen all mutants are normalized to *hcm1-3N*. Blue indicates depletion, red indicates enrichment. Shown is an average of n = 3 biological replicates. **(c)** Heat map representation of the difference between heat maps in the A/E and 3N A/E screens (from part b). Blue indicates normalized read counts were higher in 3N A/E screen, red indicates normalized read counts were higher in the A/E screen.

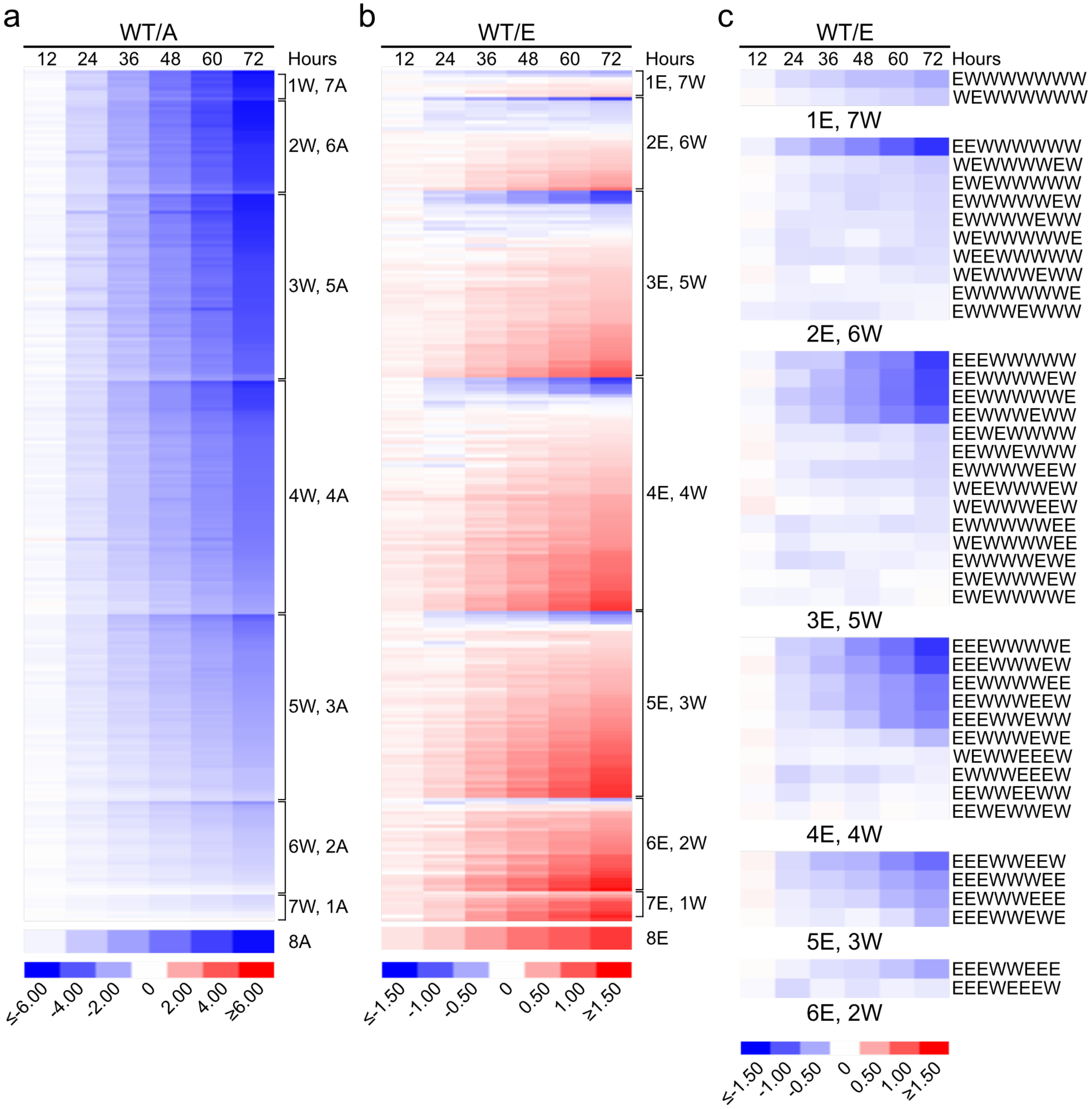


**Extended Data Fig. 5. Supporting data for the WT/A and WT/E screens. (a-b)** Heat map representation of the results of the WT/A (a) and WT/E (b) Phosphosite Scanning screens. Each row represents a mutant, shown is the log2 fold change in normalized read counts with respect to time zero for each mutant, all mutants have been normalized to wild type. Blue indicates depletion, red indicates enrichment. Shown are averages of n = 3 biological replicates. Note difference in scale bars between (a) and (b). **(c)** Expanded view of partial clusters from (b) that include mutants selected against over time. Note that all mutants in this category have phosphomimetic mutations in the first and/or second position.

### Supplementary Table 1. Strain table.

| Strain name | Genotype | Figure |
|---|---|---|
| YMC50 | *MATa his3Δ1 ura3Δ0 leu2Δ0 lys2Δ0 HIS3MX6-GAL1p-HCM1-3HA-KanMX ChrVIΔ181901-182001::Hyg-TEFp-GFP + pRS316-HCM1p-HCM1-3V5* | 1b,c, 4a |
| YMC53 | *MATa his3Δ1 ura3Δ0 leu2Δ0 lys2Δ0 HIS3MX6-GAL1p-HCM1-3HA-KanMX ChrVIΔ181901-182001::Hyg-TEFp-GFP(Y66F) + pRS316-HCM1p-HCM1-3V5* | 1b-e |
| YMC51 | *MATa his3Δ1 ura3Δ0 leu2Δ0 lys2Δ0 HIS3MX6-GAL1p-HCM1-3HA-KanMX ChrVIΔ181901-182001::Hyg-TEFp-GFP + pRS316-HCM1p-hcm1-8A-3V5* | 1d |
| YMC52 | *MATa his3Δ1 ura3Δ0 leu2Δ0 lys2Δ0 HIS3MX6-GAL1p-HCM1-3HA-KanMX ChrVIΔ181901-182001::Hyg-TEFp-GFP + pRS316-HCM1p-hcm1-8E-3V5* | 1e |
| YMC9 | *MATa his3Δ1 ura3Δ0 leu2Δ0 lys2Δ0 HIS3MX6-GAL1p-HCM1-3HA-KanMX* | 2c-g, 4e-g, 5b, c, e, f, 6a-b, E1c-d, E2, E4b-c, E5 |
| YMC390 | *MATa ade2 his3 leu2 trp1 LYS2::lexAop-HIS3 URA3::lexAop-lacZ + pBTM116* | 3a, 6c, 7c, E3 |
| YMC401 | *MATa ade2 his3 leu2 trp1 LYS2::lexAop-HIS3 URA3::lexAop-lacZ + pBTM116-hcm1(306-511)-3V5* | 3a, 6c, 7c, E3 |
| YMC402 | *MATa ade2 his3 leu2 trp1 LYS2::lexAop-HIS3 URA3::lexAop-lacZ + pBTM116-hcm1(306-511)-8A-3V5* | 3a, 6c, 7c, E3 |
| YMC403 | *MATa ade2 his3 leu2 trp1 LYS2::lexAop-HIS3 URA3::lexAop-lacZ + pBTM116-hcm1(306-511)-8E-3V5* | 3a, 6c, 7c, E3 |
| YMC395 | *MATa ade2 his3 leu2 trp1 LYS2::lexAop-HIS3 URA3::lexAop-lacZ + pBTM116-hcm1(306-511)-AAAEAAAA-3V5* | 3a, E3a |
| YMC396 | *MATa ade2 his3 leu2 trp1 LYS2::lexAop-HIS3 URA3::lexAop-lacZ + pBTM116-hcm1(306-511)-AAAAEAAA-3V5* | 3a, E3a |
| YMC397 | *MATa ade2 his3 leu2 trp1 LYS2::lexAop-HIS3 URA3::lexAop-lacZ + pBTM116-hcm1(306-511)-AAAEEAAA-3V5* | 3a, 6c, E3a-b |
| YMC398 | *MATa ade2 his3 leu2 trp1 LYS2::lexAop-HIS3 URA3::lexAop-lacZ + pBTM116-hcm1(306-511)-EEEAEEEE-3V5* | 3a, E3a |
| YMC399 | *MATa ade2 his3 leu2 trp1 LYS2::lexAop-HIS3 URA3::lexAop-lacZ + pBTM116-hcm1(306-511)-EEEEAEEE-3V5* | 3a, E3a |
| YMC400 | *MATa ade2 his3 leu2 trp1 LYS2::lexAop-HIS3 URA3::lexAop-lacZ + pBTM116-hcm1(306-511)-EEEAAEEE-3V5* | 3a, 6c, E3a-b |
| YBL192 | *MATa his3Δ1 ura3Δ0 leu2Δ0 met15Δ0 HCM1-3V5-KanMX* | 3b-d, 6d, 7b |
| YMC360 | *MATa his3Δ1 ura3Δ0 leu2Δ0 met15Δ0 hcm1-8A-3V5-KanMX* | 3b-d, 6d, 7b |
| YMC326 | *MATa his3Δ1 ura3Δ0 leu2Δ0 met15Δ0 hcm1-AAAEEAAA-3V5-KanMX* | 3b-d |
| YMC359 | *MATa his3Δ1 ura3Δ0 leu2Δ0 met15Δ0 hcm1-EEEAAEEE-3V5-KanMX* | 3b-d |
| YMC356 | *MATa his3Δ1 ura3Δ0 leu2Δ0 met15Δ0 hcm1-8E-3V5-KanMX* | 3b-d |
| YMC324 | *MATa his3Δ1 ura3Δ0 leu2Δ0 met15Δ0 hcm1-AAAEAAAA-3V5-KanMX* | 3b-d |
| YMC325 | *MATa his3Δ1 ura3Δ0 leu2Δ0 met15Δ0 hcm1-AAAAEAAA-3V5-KanMX* | 3b-d |
| YMC357 | *MATa his3Δ1 ura3Δ0 leu2Δ0 met15Δ0 hcm1-EEEAEEEE-3V5-KanMX* | 3b-d |
| YMC358 | *MATa his3Δ1 ura3Δ0 leu2Δ0 met15Δ0 hcm1-EEEEAEEE-3V5-KanMX* | 3b-d |
| YMC443 | *MATa his3Δ1 ura3Δ0 leu2Δ0 lysΔ0 HIS3MX6-GAL1p-HCM1-3HA-KanMX ChrVIΔ181901-182001::HYG-TEFp-GFP + pRS316-HCM1p-hcm1-3N-3V5* | 4b-c |

| | | |
|---|---|---|
| YMC446 | *MATa his3$\Delta$1 ura3$\Delta$0 leu2$\Delta$0 lys2$\Delta$0 HIS3MX6-GAL1p-HCM1-3HA-KanMX ChrVI$\Delta$181901-182001::Hyg-TEFp-GFP(Y66F) + pRS316-HCM1p-hcm1-3N-3V5* | 4a |
| YMC448 | *MATa his3$\Delta$1 ura3$\Delta$0 leu2$\Delta$0 lys2$\Delta$0 HIS3MX6-GAL1p-HCM1-3HA-KanMX ChrVI$\Delta$181901-182001::Hyg-TEFp-GFP(Y66F) + pRS316-HCM1p-hcm1-3N8E-3V5* | 4c |
| YMC447 | *MATa his3$\Delta$1 ura3$\Delta$0 leu2$\Delta$0 lys2$\Delta$0 HIS3MX6-GAL1p-HCM1-3HA-KanMX ChrVI$\Delta$181901-182001::Hyg-TEFp-GFP(Y66F) + pRS316-HCM1p-hcm1-3N8A-3V5* | 4b |
| YMN1 | *MATa ade2 his3 leu2 trp1 LYS2::lexAop-HIS3 URA3::lexAop-lacZ + pBTM116-hcm1(306-511)-WWWEEWWW-3V5* | 6c, E3b |
| YMN2 | *MATa ade2 his3 leu2 trp1 LYS2::lexAop-HIS3 URA3::lexAop-lacZ + pBTM116-hcm1(306-511)-EEEWWEEE-3V5* | 6c, E3b |
| YMC435 | *MATa ade2 his3 leu2 trp1 LYS2::lexAop-HIS3 URA3::lexAop-lacZ + pBTM116-hcm1(306-511)-WWWAAWWW-3V5* | 6c, E3b |
| YMC434 | *MATa ade2 his3 leu2 trp1 LYS2::lexAop-HIS3 URA3::lexAop-lacZ + pBTM116-hcm1(306-511)-AAAWWAAA-3V5* | 6c, E3b |
| YMC438 | *MATa ade2 his3 leu2 trp1 LYS2::lexAop-HIS3 URA3::lexAop-lacZ + pBTM116-hcm1(306-511)-WWWAWWWW-3V5* | 6c, E3b |
| YMC439 | *MATa ade2 his3 leu2 trp1 LYS2::lexAop-HIS3 URA3::lexAop-lacZ + pBTM116-hcm1(306-511)-WWWWAWWW-3V5* | 6c, E3b |
| YBL193 | *MATa his3$\Delta$1 ura3$\Delta$0 leu2$\Delta$0 met15$\Delta$0 hcm1-15A-3V5-KanmX* | 6d, 7b |
| YMC426 | *MATa his3$\Delta$1 ura3$\Delta$0 leu2$\Delta$0 met15$\Delta$0 hcm1-AAAWAAAA-3V5-KanMX* | 6d |
| YMC427 | *MATa his3$\Delta$1 ura3$\Delta$0 leu2$\Delta$0 met15$\Delta$0 hcm1-AAAAWAAA-3V5-KanMX* | 6d |
| YMC423 | *MATa his3$\Delta$1 ura3$\Delta$0 leu2$\Delta$0 met15$\Delta$0 hcm1-AAAWWAAA-3V5-KanMX* | 6d, 7b |
| YMC425 | *MATa his3$\Delta$1 ura3$\Delta$0 leu2$\Delta$0 met15$\Delta$0 hcm1-WWWAAWWW-3V5-KanMX* | 6d |
| YMC428 | *MATa his3$\Delta$1 ura3$\Delta$0 leu2$\Delta$0 met15$\Delta$0 hcm1-WWWAWWWW-3V5-KanMX* | 6d |
| YMC429 | *MATa his3$\Delta$1 ura3$\Delta$0 leu2$\Delta$0 met15$\Delta$0 hcm1-WWWWAWWW-3V5-KanMX* | 6d |
| YMC430 | *MATa his3$\Delta$1 ura3$\Delta$0 leu2$\Delta$0 met15$\Delta$0 hcm1-SSSSWWWW-3V5-KanMX* | 7b |
| YMC431 | *MATa his3$\Delta$1 ura3$\Delta$0 leu2$\Delta$0 met15$\Delta$0 hcm1-SSSWWWWW-3V5-KanMX* | 7b |
| YMC432 | *MATa his3$\Delta$1 ura3$\Delta$0 leu2$\Delta$0 met15$\Delta$0 hcm1-WWWSWWWW-3V5-KanMX* | 7b |
| YMC440 | *MATa ade2 his3 leu2 trp1 LYS2::lexAop-HIS3 URA3::lexAop-lacZ + pBTM116-hcm1(306-511)-SSSSWWWW-3V5* | 7c, E3c |
| YMC441 | *MATa ade2 his3 leu2 trp1 LYS2::lexAop-HIS3 URA3::lexAop-lacZ + pBTM116-hcm1(306-511)-SSSWWWWW-3V5* | 7c, E3c |
| YMC442 | *MATa ade2 his3 leu2 trp1 LYS2::lexAop-HIS3 URA3::lexAop-lacZ + pBTM116-hcm1(306-511)-WWWSWWWW-3V5* | 7c, E3c |
| YMC343 | *MATa his3$\Delta$1 ura3$\Delta$0 leu2$\Delta$0 met15$\Delta$0 + pRS316-HCM1p-HCM1-3V5* | E1b |
| YMC344 | *MATa his3$\Delta$1 ura3$\Delta$0 leu2$\Delta$0 met15$\Delta$0 + pRS316-HCM1p-hcm1-8A-3V5* | E1b |
| YMC345 | *MATa his3$\Delta$1 ura3$\Delta$0 leu2$\Delta$0 met15$\Delta$0 + pRS316-HCM1p-hcm1-8E-3V5* | E1b |
| YMC346 | *MATa his3$\Delta$1 ura3$\Delta$0 leu2$\Delta$0 met15$\Delta$0 + pRS316-HCM1p-hcm1-AAEEAEAE-3V5* | E1b |
| YMC347 | *MATa his3$\Delta$0 ura3$\Delta$0 leu2$\Delta$0 met15$\Delta$0 + pRS316-HCM1p-hcm1-EAAAAAAE-3V5* | E1b |
| YMC348 | *MATa his3$\Delta$1 ura3$\Delta$0 leu2$\Delta$0 met15$\Delta$0 + pRS316-HCM1p-hcm1-AAEAAAEE-3V5* | E1b |

| | | |
|---|---|---|
| YMC349 | *MATa his3Δ1 ura3Δ0 leu2Δ0 met15Δ0 + pRS316-HCM1p-hcm1-AAEAAAAA-3V5* | E1b |
| YMC350 | *MATa his3Δ1 ura3Δ0 leu2Δ0 met15Δ0 + pRS316-HCM1p-hcm1-AAAEEAAE-3V5* | E1b |
| YMC351 | *MATa his3Δ1 ura3Δ0 leu2Δ0 met15Δ0 + pRS316-HCM1p-hcm1-EEAAAEAE-3V5* | E1b |
| YMC352 | *MATa his3Δ1 ura3Δ0 leu2Δ0 met15Δ0 + pRS316-HCM1p-hcm1-AEEAEEEE-3V5* | E1b |
| YMC353 | *MATa his3Δ0 ura3Δ0 leu2Δ0 met15Δ0 + pRS316-HCM1p-hcm1-EEEEAEAE-3V5* | E1b |
| YMC354 | *MATa his3Δ1 ura3Δ0 leu2Δ0 met15Δ0 + pRS316-HCM1p-hcm1-EAAAEAAA-3V5* | E1b |
| YMC355 | *MATa his3Δ1 ura3Δ0 leu2Δ0 met15Δ0 + pRS316-HCM1p-hcm1-AAEAEEAE-3V5* | E1b |
| YMC454 | *MATa his3Δ1 ura3Δ0 leu2Δ0 met15Δ0 hcm1Δ::KanMX + pRS316* | E4a |
| YMC455 | *MATa his3Δ1 ura3Δ0 leu2Δ0 met15Δ0 hcm1Δ::KanMX + pRS316-HCM1-3V5* | E4a |
| YMC458 | *MATa his3Δ1 ura3Δ0 leu2Δ0 met15Δ0 hcm1Δ::KanMX + pRS316-hcm1-3N-3V5* | E4a |

**Supplementary Table 2. Plasmid table.**

| Plasmid name | Description | Figure |
|---|---|---|
| pRS316-HCM1-3V5 | *HCM1p-HCM1-3V5*, CEN, URA3 | 1b-e, 4a, E4a, E1b |
| pRS316-hcm1-8A-3V5 | *HCM1p-hcm1-8A-3V5*, CEN, URA3 | 1d, E1b |
| pRS316-hcm1-8E-3V5 | *HCM1p-hcm1-8E-3V5*, CEN, URA3 | 1e, E1b |
| pRS316-hcm1-3N-3V5 | *HCM1p-hcm1-3N-3V5,* CEN, URA3 | 4a-c, E4a |
| pRS316-hcm1-3N8A-3V5 | *HCM1p-hcm1-3N8A-3V5*, CEN, URA3 | 4b |
| pRS316-hcm1-3N8E-3V5 | *HCM1p-hcm1-3N8E-3V5*, CEN, URA3 | 4c |
| pBTM116 | *ADH1p-LEXA*, 2micron, TRP | 3a, 6c, 7c, E3a-c |
| pBTM116-hcm1(306-511)-3V5 | *ADH1p-LEXA-hcm1(306-511)-3V5*, 2micron, TRP | 3a, 6c, 7c, E3a-c |
| pBTM116-hcm1(306-511)-8A-3V5 | *ADH1p-LEXA-hcm1(306-511)-8A-3V5*, 2micron, TRP | 3a, 6c, 7c, E3a-c |
| pBTM116-hcm1(306-511)-AAAEAAAA-3V5 | *ADH1p-LEXA-hcm1(306-511)-AAAEAAAA-3V5*, 2micron, TRP | 3a, E3a |
| pBTM116-hcm1(306-511)-AAAAEAAA-3V5 | *ADH1p-LEXA-hcm1(306-511)-AAAAEAAA-3V5*, 2micron, TRP | 3a, E3a |
| pBTM116-hcm1(306-511)-AAAEEAAA-3V5 | *ADH1p-LEXA-hcm1(306-511)-AAAEEAAA-3V5*, 2micron, TRP | 3a, 6c, E3a-b |
| pBTM116-hcm1(306-511)-EEEAEEEE-3V5 | *ADH1p-LEXA-hcm1(306-511)-EEEAEEEE-3V5*, 2micron, TRP | 3a, E3a |
| pBTM116-hcm1(306-511)-EEEEAEEE-3V5 | *ADH1p-LEXA-hcm1(306-511)-EEEEAEEE-3V5*, 2micron, TRP | 3a, E3a |
| pBTM116-hcm1(306-511)-EEEAAEEE-3V5 | *ADH1p-LEXA-hcm1(306-511)-EEEAAEEE-3V5*, 2micron, TRP | 3a, 6c, E3a-b |
| pBTM116-hcm1(306-511)-8E-3V5 | *ADH1p-LEXA-hcm1(306-511)-8E-3V5*, 2micron, TRP | 3a, 6c, 7c, E3a-c |
| pBTM116-hcm1(306-511)-AAAWWAAA-3V5 | *ADH1p-LEXA-hcm1(306-511)-AAAWWAAA-3V5*, 2micron, TRP | 6c, E3b |
| pBTM116-hcm1(306-511)-WWWAAWWW-3V5 | *ADH1p-LEXA-hcm1(306-511)-WWWAAWWW-3V5*, 2micron, TRP | 6c, E3b |
| pBTM116-hcm1(306-511)-WWWAWWWW-3V5 | ADH1p-LEXA-hcm1(306-511)-WWWAWWWW-3V5, 2micron, TRP | 6c, E3b |
| pBTM116-hcm1(306-511)-WWWWAWWW-3V5 | *ADH1p-LEXA-hcm1(306-511)-WWWWAWWW-3V5*, 2micron, TRP | 6c, E3b |
| pBTM116-hcm1(306-511)-WWWEEWWW-3V5 | *ADH1p-LEXA-hcm1(306-511)-WWWEEWWW-3V5*, 2micron, TRP | 6c, E3b |
| pBTM116-hcm1(306-511)-EEEWWEEE-3V5 | *ADH1p-LEXA-hcm1(306-511)-EEEWWEEE-3V5*, 2micron, TRP | 6c, E3b |
| pBTM116-hcm1(306-511)-SSSSWWWW-3V5 | *ADH1p-LEXA-hcm1(306-511)-SSSSWWWW-3V5*, 2micron, TRP | 7c, E3c |
| pBTM116-hcm1(306-511)-SSSWWWWW-3V5 | *ADH1p-LEXA-hcm1(306-511)-SSSWWWWW-3V5*, 2micron, TRP | 7c, E3c |
| pBTM116-hcm1(306-511)-WWWSWWWW-3V5 | *ADH1p-LEXA-hcm1(306-511)-WWWSWWWW-3V5*, 2micron, TRP | 7c, E3c |
| pRS316-hcm1-AAEEAEAE-3V5 | *HCM1p-hcm1-AAEEAEAE-3V5*, CEN, URA3 | E1b |
| pRS316-hcm1-EAAAAAAE-3V5 | *HCM1p-hcm1-EAAAAAAE-3V5*, CEN, URA3 | E1b |

| | | |
|---|---|---|
| pRS316-hcm1-AAEAAAEE-3V5 | *HCM1p-hcm1-AAEAAAEE-3V5*, CEN, URA3 | E1b |
| pRS316-hcm1-AAEAAAAA-3V5 | *HCM1p-hcm1-AAEAAAAA-3V5*, CEN, URA3 | E1b |
| pRS316-hcm1-AAAEEAAE-3V5 | *HCM1p-hcm1-AAAEEAAE-3V5*, CEN, URA3 | E1b |
| pRS316-hcm1-EEAAAEAE-3V5 | *HCM1p-hcm1-EEAAAEAE-3V5*, CEN, URA3 | E1b |
| pRS316-hcm1-AEEAEEEE-3V5 | *HCM1p-hcm1-AEEAEEEE-3V5*, CEN, URA3 | E1b |
| pRS316-hcm1-EEEEAEAE-3V5 | *HCM1p-hcm1-EEEEAEAE-3V5*, CEN, URA3 | E1b |
| pRS316-hcm1-EAAAEAAA-3V5 | *HCM1p-hcm1-EAAAEAAA-3V5*, CEN, URA3 | E1b |
| pRS316-hcm1-AAEAEEAE-3V5 | *HCM1p-hcm1-AAEAEEAE-3V5*, CEN, URA3 | E1b |
| pRS316-hcm1-A/E | *HCM1p-hcm1-A/E-3V5,* CEN, URA3 | 2c-g, 4g, 6a-b, S1c-d, E2a, E4b-c |
| pRS316-hcm1-3NA/E | *HCM1p-hcm1-3NA/E-3V5,* CEN, URA3 | 4e-g, E2b, E4b-c |
| pRS316-hcm1-WT/A | *HCM1p-hcm1-WT/A-3V5,* CEN, URA3 | 5b-c, 6b, E2c, E5a |
| pRS316-hcm1-WT/E | *HCM1p-hcm1-WT/E-3V5,* CEN, URA3 | 5e-f, 6a-b, E2d, E5b-c |

### Supplementary Table 3. Oligonucleotide table

| Name | Sites included | Genotype | Oligo orientation | Sequence |
|---|---|---|---|---|
| A1 | T428 | A | FWD | ccttcctctcatggttcggacttacttaaa**GCTCCT**aagatgaggcattccgatggcttagagaaa |
| A2 | T428 | E | FWD | ccttcctctcatggttcggacttacttaaa**GAGGAA**aagatgaggcattccgatggcttagagaaa |
| A3 | T428 | W | FWD | ccttcctctcatggttcggacttacttaaa**ACACCC**aagatgaggcattccgatggcttagagaaa |
| B1 | T440, T447 | AA | REV | ctgccatttcctcaaaatcgagttaccgtcctt**TGGCGC**gcttatcaaccgcga**CGGAGC**tttctctaagccatcggaatgc |
| B2 | T440, T447 | AE | REV | ctgccatttcctcaaaatcgagttaccgtcctt**CTCTTC**gcttatcaaccgcga**CGGAGC**tttctctaagccatcggaatgc |
| B3 | T440, T447 | EA | REV | ctgccatttcctcaaaatcgagttaccgtcctt**TGGCGC**gcttatcaaccgcga**TTCCTC**tttctctaagccatcggaatgc |
| B4 | T440, T447 | EE | REV | ctgccatttcctcaaaatcgagttaccgtcctt**CTCTTC**gcttatcaaccgcga**TTCCTC**tttctctaagccatcggaatgc |
| B5 | T440, T447 | AW | REV | ctgccatttcctcaaaatcgagttaccgtcctt**AGGTGT**gcttatcaaccgcga**CGGAGC**tttctctaagccatcggaatgc |
| B6 | T440, T447 | WA | REV | ctgccatttcctcaaaatcgagttaccgtcctt**TGGCGC**gcttatcaaccgcga**TGGGGT**tttctctaagccatcggaatgc |
| B7 | T440, T447 | WW | REV | ctgccatttcctcaaaatcgagttaccgtcctt**AGGTGT**gcttatcaaccgcga**TGGGGT**tttctctaagccatcggaatgc |
| B8 | T440, T447 | WE | REV | ctgccatttcctcaaaatcgagttaccgtcctt**CTCTTC**gcttatcaaccgcga**TGGGGT**tttctctaagccatcggaatgc |
| B9 | T440, T447 | EW | REV | ctgccatttcctcaaaatcgagttaccgtcctt**AGGTGT**gcttatcaaccgcga**TTCCTC**tttctctaagccatcggaatgc |
| C1 | T460 | A | FWD | actcgattttgaggaaatggcag**GCACCA**tcacaccttttgaagatttgtactgt |
| C2 | T460 | E | FWD | actcgattttgaggaaatggcag**GAGGAG**tcacaccttttgaagatttgtactgt |
| C3 | T460 | W | FWD | actcgattttgaggaaatggcag**ACTCCT**tcacaccttttgaagatttgtactgt |
| D1 | S471, T479, T486 | AAA | REV | aatttgggtttccaaagtgccccc**GGGGGC**cgtgatatacctgat**AGGAGC**ctctatagctctaaatag**GGGTGC**acagtacaaatcttcaaaaaggtgtga |
| D2 | S471, T479, T486 | AAE | REV | aatttgggtttccaaagtgccccc**TTCCTC**cgtgatatacctgat**AGGAGC**ctctatagctctaaatag**GGGTGC**acagtacaaatcttcaaaaaggtgtga |
| D3 | S471, T479, T486 | AEA | REV | aatttgggtttccaaagtgccccc**GGGGGC**cgtgatatacctgat**CTCTTC**ctctatagctctaaatag**GGGTGC**acagtacaaatcttcaaaaaggtgtga |
| D4 | S471, T479, T486 | AEE | REV | aatttgggtttccaaagtgccccc**TTCCTC**cgtgatatacctgat**CTCTTC**ctctatagctctaaatag**GGGTGC**acagtacaaatcttcaaaaaggtgtga |
| D5 | S471, T479, T486 | EAA | REV | aatttgggtttccaaagtgccccc**GGGGGC**cgtgatatacctgat**AGGAGC**ctctatagctctaaatag**TTCCTC**acagtacaaatcttcaaaaaggtgtga |
| D6 | S471, T479, T486 | EAE | REV | aatttgggtttccaaagtgccccc**TTCCTC**cgtgatatacctgat**AGGAGC**ctctatagctctaaatag**TTCCTC**acagtacaaatcttcaaaaaggtgtga |
| D7 | S471, T479, T486 | EEA | REV | aatttgggtttccaaagtgccccc**GGGGGC**cgtgatatacctgat**CTCTTC**ctctatagctctaaatag**TTCCTC**acagtacaaatcttcaaaaaggtgtga |

| | | | | |
|---|---|---|---|---|
| D8 | S471, T479, T486 | EEE | REV | aatttgggtttccaaagtgcccccTTCCTCcgtgatatacctgatCTCTTCctctatagctctaaatagTTCCTCacagtacaaatcttcaaaaaggtgtga |
| D9 | S471, T479, T486 | AAW | REV | aatttgggtttccaaagtgcccccCGGCGTcgtgatatacctgatAGGAGCctctatagctctaaatagGGGTGCacagtacaaatcttcaaaaaggtgtga |
| D10 | S471, T479, T486 | AWA | REV | aatttgggtttccaaagtgcccccGGGGGCcgtgatatacctgatTGGAGTctctatagctctaaatagGGGTGCacagtacaaatcttcaaaaaggtgtga |
| D11 | S471, T479, T486 | AWW | REV | aatttgggtttccaaagtgcccccCGGCGTcgtgatatacctgatTGGAGTctctatagctctaaatagGGGTGCacagtacaaatcttcaaaaaggtgtga |
| D12 | S471, T479, T486 | WAA | REV | aatttgggtttccaaagtgcccccGGGGGCcgtgatatacctgatAGGAGCctctatagctctaaatagCGGAGAacagtacaaatcttcaaaaaggtgtga |
| D13 | S471, T479, T486 | WAW | REV | aatttgggtttccaaagtgcccccCGGCGTcgtgatatacctgatAGGAGCctctatagctctaaatagCGGAGAacagtacaaatcttcaaaaaggtgtga |
| D14 | S471, T479, T486 | WWA | REV | aatttgggtttccaaagtgcccccGGGGGCcgtgatatacctgatTGGAGTctctatagctctaaatagCGGAGAacagtacaaatcttcaaaaaggtgtga |
| D15 | S471, T479, T486 | WWW | REV | aatttgggtttccaaagtgcccccCGGCGTcgtgatatacctgatTGGAGTctctatagctctaaatagCGGAGAacagtacaaatcttcaaaaaggtgtga |
| D16 | S471, T479, T486 | WWE | REV | aatttgggtttccaaagtgcccccTTCCTCcgtgatatacctgatTGGAGTctctatagctctaaatagCGGAGAacagtacaaatcttcaaaaaggtgtga |
| D17 | S471, T479, T486 | WEW | REV | aatttgggtttccaaagtgcccccCGGCGTcgtgatatacctgatCTCTTCctctatagctctaaatagCGGAGAacagtacaaatcttcaaaaaggtgtga |
| D18 | S471, T479, T486 | WEE | REV | aatttgggtttccaaagtgcccccTTCCTCcgtgatatacctgatCTCTTCctctatagctctaaatagCGGAGAacagtacaaatcttcaaaaaggtgtga |
| D19 | S471, T479, T486 | EWW | REV | aatttgggtttccaaagtgcccccCGGCGTcgtgatatacctgatTGGAGTctctatagctctaaatagTTCCTCacagtacaaatcttcaaaaaggtgtga |
| D20 | S471, T479, T486 | EWE | REV | aatttgggtttccaaagtgcccccTTCCTCcgtgatatacctgatTGGAGTctctatagctctaaatagTTCCTCacagtacaaatcttcaaaaaggtgtga |
| D21 | S471, T479, T486 | EEW | REV | aatttgggtttccaaagtgcccccCGGCGTcgtgatatacctgatCTCTTCctctatagctctaaatagTTCCTCacagtacaaatcttcaaaaaggtgtga |
| E1 | S496 | A | FWD | ggcactttggaaacccaaattGCTCCTagaaagtcctctgcacccgat |
| E2 | S496 | E | FWD | ggcactttggaaacccaaattGAGGAGagaaagtcctctgcacccgat |
| E3 | S496 | W | FWD | ggcactttggaaacccaaattTCACCAagaaagtcctctgcacccgat |
| F1 | N/A | N/A | REV | tgtgaggacatcgggtgcagaggactttct |

**Supplementary File 1.**

```
import itertools, re, gzip, os, sys, warnings
import matplotlib; matplotlib.use('agg')
import matplotlib.pyplot as plt;
from pathlib import Path


def reverse_complement(dna):
    complement = {'A': 'T', 'C': 'G', 'G': 'C', 'T': 'A', 'N':'N'}
    return ''.join([complement[base] for base in dna[::-1]])


# Read CMD
fname1 = sys.argv[1]
fname2 = sys.argv[2]
name = sys.argv[3]
print("CMD: ", sys.argv)

# Mkdir
Path("pie1").mkdir(parents=True, exist_ok=True)
Path("pie2").mkdir(parents=True, exist_ok=True)
Path("count").mkdir(parents=True, exist_ok=True)


# Params
## C: Common Seq
## S: Site Seq
C0 = "CTCATGGTTCGGACTTACTTAAA"
C1 = "AAGATGAGGCATTCCGATGGCTTAGAGAAA"
C2 = "TCGCGGTTGATAAGC"
C3 = "AAGGACGGTAACTCGATTTTGAGGAAATGGCAG"
C4 = "TCACACCTTTTTGAAGATTTGTACTGT"
C5 = "CTATTTAGAGCTATAGAG"
C6 = "ATCAGGTATATCACG"
C7 = "GGGGGCACTTTGGAAACCCAAATT"
C8 = "AGAAAGTCCTCTGCACCC"
Commons = [C0, C1, C2, C3, C4, C5, C6, C7, C8]

C = {}
for i in range(0, 9):
    C["C{0}".format(i)] = Commons[i]
print(C)

S0 = ["GCTCCT", "GAGGAA"]
S1 = ["GCTCCG", "GAGGAA"]
S2 = ["GCGCCA", "GAAGAG"]
S3 = ["GCACCA", "GAGGAG"]
S4 = ["GCACCC", "GAGGAA"]
S5 = ["GCTCCT", "GAAGAG"]
S6 = ["GCCCCC", "GAGGAA"]
S7 = ["GCTCCT", "GAGGAG"]
Sites = [S0, S1, S2, S3, S4, S5, S6, S7]
```

```
S = {}
for i in range(0,8):
    S["S{0}".format(i)] = Sites[i]
print(S)


# Get MutSeq
combinations = list(itertools.product([0,1], repeat=8)) # 0-255
print(combinations[0],
combinations[1],
combinations[255])

dictionary = dict(zip([0,1], ["A", "E"]))
dictionary

mutSeqDict = {}
for i in range(0, 256):
    #print("\n", i)

    combination = combinations[i]
    #print(combination)

    letter_lst = [dictionary[x] for x in combination]
    letter = "".join(letter_lst)
    #print(">{0}".format(letter))

    site_seqs = []
    for j in range(0, 8):
        Skey = "S{0}".format(j)
        seq = S[Skey]
        #print(Skey, seq)

        select = combination[j]
        #print(Skey, seq[select])
        site_seqs.append(seq[select])

    common_seqs = Commons

    mut_seqs = list(zip(common_seqs, site_seqs))
    mut_seqs.append(common_seqs[8])
    mut_seqs
    join1 = [''.join(x) for x in mut_seqs]
    join2 = "".join(join1)
    mutSeq = join2
    # print(mutSeq)

    mutSeqDict[letter] = mutSeq

mutSeqDict["WWWWWWWW"] =
"CTCATGGTTCGGACTTACTTAAAACACCCAAGATGAGGCATTCCGATGGCTTAGAGAAAACCCCATCGCGGT
TGATAAGCACACCTAAGGACGGTAACTCGATTTTGAGGAAATGGCAGACTCCTTCACACCTTTTTGAAGATTT
GTACTGTTCTCCGCTATTTAGAGCTATAGAGACTCCAATCAGGTATATCACGACGCCGGGGGGCACTTTGGAA
ACCCAAATTTCACCAAGAAAGTCCTCTGCACCC"
```

```
print(len(mutSeqDict), " Mut seqs")


# Save Mut.Fasta
file = open("MutSeqs257.fasta", 'w')
for key in mutSeqDict:
    seq = mutSeqDict[key]
    file.writelines(">{0}\n{1}\n".format(key, seq))
file.close()


# Read Fastq and Exact Match
def exact_match(fname1, fname2, mutSeqDict=mutSeqDict, test = False):
    count_dict = {}
    n = 0
    with gzip.open(fname1, 'rt') as R1, gzip.open(fname2, 'rt') as R2: #
read PE data
        for line1, line2 in zip(R1, R2):
            n += 1
            line1 = line1.strip()
            line2 = line2.strip()

            if n % 4 == 2: # for seq line
                #print("{0}\n{1}\n".format(line1, line2))
                read1 = line1#[5:145] # params no trimming
                read2 = line2#[5:145] # params no trimming
                #read1 = line1[24:86] # test
                #read2 = line2[17:60] # test
                read2_rc = reverse_complement(read2)
                #print("{0}\n{1}\n{2}\n".format(read1, read2, read2_rc))

                match_flag = 0
                mem_key = ""
                for key in mutSeqDict:
                    mutseq = mutSeqDict[key] # not simply mapping, length
issue, 251 bp

                    if re.search(read1, mutseq): # R1 search
                        if re.search(read2_rc, mutseq): # R2 search
                            #print(n, "read1 both match", key, read1,
mutseq)
                            #print(n, "read2 both match", key, read2,
mutseq)
                            #print(key)
                            count_dict[key] = count_dict.get(key, 0) + 1
# .get allows you to specify a default value if the key does not exist.
                            if match_flag: # Bug, if match more than 2
patterns (because of testing shortcuts, should not have this in
production code)
                                warnings.warn("More than two matches
found for fastq line{0}: {1}, {2}".format(n, mem_key, key))
                            mem_key = key
                            match_flag += 1
```

```
            if test:
                if n > 120000:
                    break
    return [count_dict, n//4]


# Exact Match and Count
[count_dict, total] = exact_match(fname1, fname2, test = 0)

# Pie plot discarding 'Other' mutations
labels = list(count_dict.keys())
sizes = list(count_dict.values())

fig1, ax1 = plt.subplots()
ax1.pie(sizes,
        labels=labels,
        autopct='%1.1f%%',
        shadow=True, startangle=90)
ax1.axis('equal')  # Equal aspect ratio ensures that pie is drawn as a
circle.
plt.savefig("pie1/" + name + ".all.pdf")

# Pie plot including 'Other' mutations
labels = list(count_dict.keys())
sizes = list(count_dict.values())
size_other = total - sum(sizes)
labels.append("other")
sizes.append(size_other)

fig1, ax1 = plt.subplots()
ax1.pie(sizes,
        labels=labels,
        autopct='%1.1f%%',
        shadow=True, startangle=90)
ax1.axis('equal')  # Equal aspect ratio ensures that pie is drawn as a
circle.
plt.savefig("pie2/" + name + ".other.pdf")

# Save

file = open("count/"+name+".txt", 'w')
file.writelines("# ID,count\n")
for key in count_dict:
    count = count_dict[key]
    file.writelines("{0}, {1}\n".format(key, count))
file.writelines("Others,{0}\n".format(size_other))
file.close()
```
